# Apoptosis-induced self-renewal of neurogenic progenitors safeguards retinal development against extensive cell loss

**DOI:** 10.64898/2026.07.03.735591

**Authors:** Catarina Figueiredo, Greta Tellkamp, Caren Norden, Mauricio Rocha-Martins

## Abstract

Developing embryos have a striking ability to buffer external and internal perturbations. A key example of this phenomenon across developing systems is successful organ formation even after substantial cell loss. Although molecular regulators of this robustness are beginning to be understood, it remains unclear whether and how surviving cells can remodel their developmental trajectories to safeguard morphogenesis. To address this question, we use the zebrafish retina, where progenitor behaviour and lineage transitions are highly tractable and extensively characterized. We induce widespread apoptosis via heat stress and genetic approaches and track cellular and tissue-wide responses *in vivo* over time. We find that retinal development is highly resilient with neurogenesis initiating on time and growth continuing despite extensive apoptosis. Continued growth is supported by neurogenic progenitors that react to apoptosis through a non-cell autonomous switch in behaviour. These cells bypass their canonical differentiation route and undergo self-renewing divisions that expand clonal output and compensate for lost cells. Importantly, self-renewal is transient, progenitors resume lineage progression, generating appropriate neuronal cell types. This adaptive response supports the formation of retinas with proper architecture, connectivity to the brain and visual function. Together, these findings identify latent plasticity in the neurogenic programme as a mechanism that contributes to developmental robustness under stress.

## Introduction

Developing embryos have a remarkable capacity to generate functional organs reproducibly even when challenged by internal or external stressors^1^. External stressors vary depending on organism and context and include variations in temperature, hypoxia, pathogen exposure and UV radiation^2^. The possibility of embryos to react to these challenges implies that development features buffering capacities that allow the organism to resist not only genetic variation but also environmental insults^3,4^. If such buffering mechanisms fail, developmental stress can have severe consequences. For example, failed adaptation can impair cell proliferation, disrupt cell movements or even trigger cell death which, in the worst cases, can compromise organ and organismal formation^5–7^. Thus, the understanding of embryonic adaptation to stress is crucial to explain how reproducible embryogenesis is achieved.

Some of the molecular mechanisms that provide resilience to cellular stressors have been identified^8–10^. These include protein quality control mediated by chaperones that has been shown to prevent cytotoxicity caused by the accumulation of misfolded proteins^11–13^. Another example is the *Drosophila* gap gene system, where gene regulatory networks can buffer morphogen noise and ensure reproducible patterning despite deleterious mutations, thereby preventing cellular defects^14,15^. At the cellular and tissue level, embryos can adapt developmental trajectories and remodel tissues to buffer or correct for errors^16–18^. This buffering capacity is seen by the ability of developing tissues to recover growth after substantial cell loss. Compensatory proliferation in response to acute perturbations that cause cell loss is seen in the *Drosophila* wing disc and eye^19–21^ and the mouse cortex^22^. Studies in *Drosophila* revealed that caspase activation in dying cells can promote the secretion of mitogens that induce proliferation of surviving neighbours, indicating that compensatory growth is a direct response to cell death^23,24^. However, most of these studies have largely relied on static snapshots and endpoint analyses, leaving the dynamics of tissue-wide adaptations and cellular plasticity during growth recovery underexplored.

In the developing central nervous system (CNS) robustness to cell loss is particularly critical as cell number, tissue growth and neural connectivity need to be tightly regulated and unwanted loss of cells can have direct functional consequences^25,26^. This makes the CNS an attractive model to study whether and how surviving cells adjust their developmental programme to ensure proper organ formation. During normal CNS development, low levels of programmed cell death can occur to eliminate defective cells^27–29^. However, stressors including heat stress, hypoxia, viral infections and genetic abnormalities can markedly increase cell death. For example, genetic mutations affecting the replication stress response or ZIKV viral infections can cause neural progenitor death and lead to microcephaly and retinal malformations^30,31^. Despite these abundant stressors that can perturb neural development, overt developmental defects are relatively rare in humans (27.55 per 10000 births in the EU; (EUROCAT, 2026)^32^). Thus, it has been suggested that neural tissues can adapt to cell loss to ensure proper organ development^33,34^. Consistent with this idea, chicken embryos can fully recover from partial ablation of neural tube segments^35,36^. Similarly, studies of human brain organoids and zebrafish retina indicate that the induction of mosaic apoptosis by selective elimination of cells can be counteracted by neural progenitors adjusting their clone size^37,38^. These findings suggest that neural progenitors have an intrinsic capacity to sense and compensate for cell loss, thereby safeguarding tissue growth. One source of this flexibility could stem from the fact that individual progenitor decisions are largely stochastic in brain and retinal tissue. This has been shown to result in variable clone sizes^39–41^. However, whether this variability could also provide a source of adaptive responses to stress and cell loss remains currently unclear.

We here use the zebrafish retina, in which progenitor division patterns and lineage transitions have been characterized^40,42–45^, to investigate robustness against elevated cell death. We induced high levels of transient apoptosis through different experimental regimes and found that rather than stalling development, retinas experiencing extensive cell death initiate neurogenesis at the correct developmental stage and, at the same time, continue to grow. This continued growth is driven by a non-cell-autonomous response in which a major population of neurogenic progenitors (Ath5+) bypasses the canonical transition to differentiating cell states leading to self-renewing divisions. These additional divisions expand the progenitor pool and thereby compensate for cell loss. Self-renewing neurogenic progenitors ultimately resume normal developmental trajectories and contribute to the generation of functional retinas with appropriate architecture, connectivity to the brain and visual function. This indicates that plasticity within the neurogenic program has the capacity to buffer retinal formation against cell loss during differentiation.

## Results

### Retinal development progresses despite extensive heat stress-induced apoptosis

Previous studies have shown that the developing central nervous system (CNS) is sensitive to apoptosis upon heat stress as seen in the brains of rodents^46,47^, lizards^48^ and zebrafish^49^. We therefore used global heat stress to induce widespread cell death in the developing CNS of zebrafish at early stages of organogenesis. To this end, 24 hours post-fertilization (hpf) embryos were exposed to 42 °C for 45 minutes (Figure 1A). Staining against the pro-apoptotic marker active caspase 3 one day after heat stress (48 hpf) revealed that indeed apoptosis occurred particularly prominently in the CNS, including the retina (Figure 1B) in all embryos albeit to varying degrees of severity (Figure 1B, 1C, Figure S1B). Despite this extensive cell death at 48 hpf, one day after heat stress, 94% of embryos had survived (Figure S1A) and 83% of embryos survived until 72 hpf (Figure S1A).

**Figure 1.**
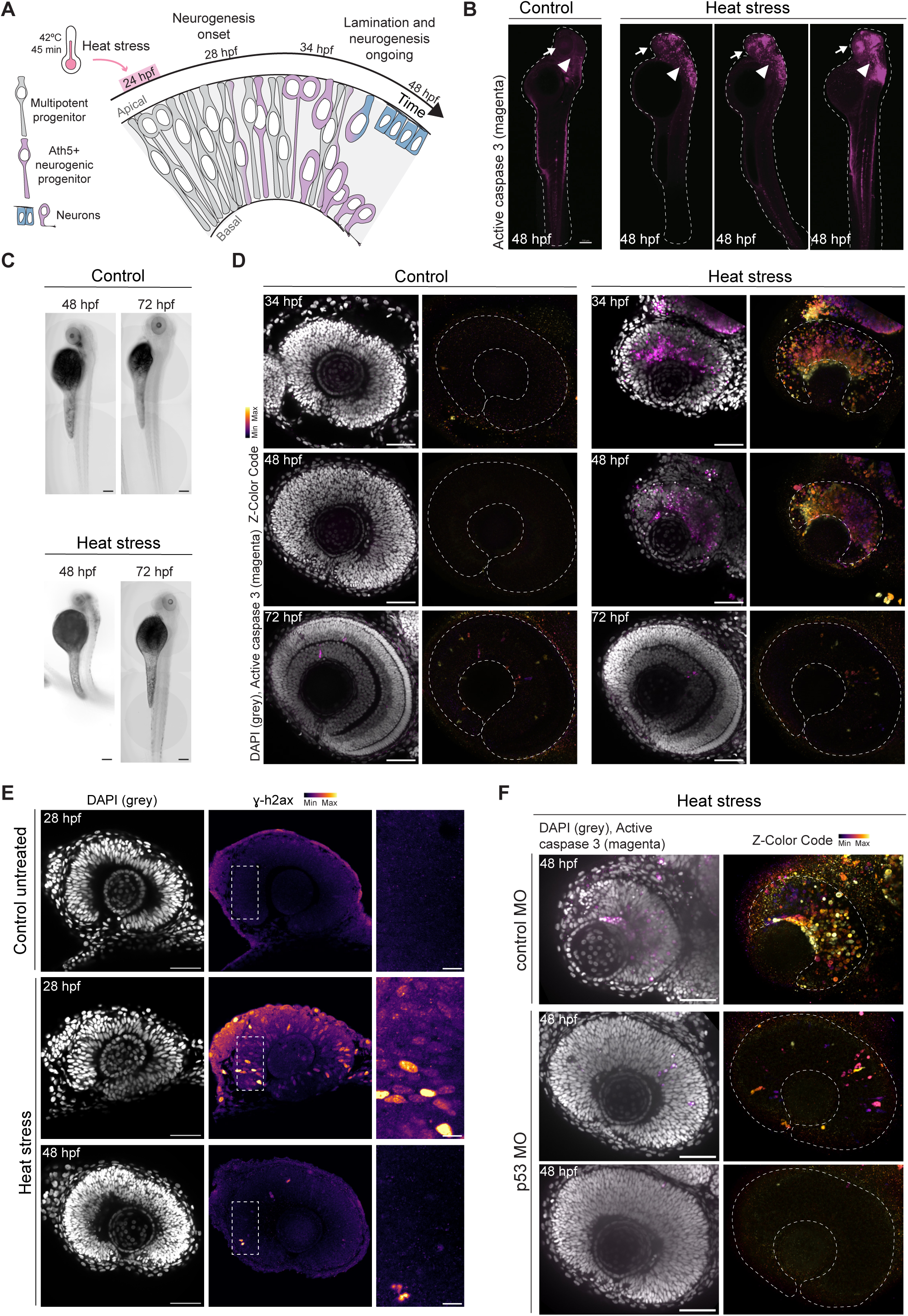
Heat stress induces transient apoptosis in the developing zebrafish retina. **(A)** Schematic of retinal development following heat stress. At 24 hpf, when embryos are subject to heat stress, the retina is populated only by multipotent progenitors. By 28 hpf, neurogenesis starts as progenitors begin expressing Ath5 and by 48 hpf most neurons are differentiated and organized in distinct layers. **(B)** Spatial distribution of apoptotic cells in whole embryos at 48 hpf. Active caspase 3 (magenta) labels apoptotic cells. Arrows and arrowheads indicate apoptotic cells in the retina and in the brain, respectively. **(C)** Brightfield images of control and heat-stressed embryos at 48 hpf and 72 hpf. **(D)** Spatial distribution of apoptotic cells in retinas of control (left) and heat-stressed (right) embryos at 10 h (34 hpf; top), 24 h (48 hpf; middle) and 48 h (72 hpf; bottom) after heat stress. DAPI (grey) labels nuclei. Active caspase 3 labels apoptotic cells; colour represents Z depth. **(E)** Increased replicative stress in heat-stressed retinas. Control (top) and heat-stressed retinal neuroepithelia at 4 h (middle) and 24 h (bottom) after heat stress. DAPI (grey) labels nuclei and ɣh2ax labels DNA double-strand breaks; colour represents fluorescence intensity. Dashed white boxes indicate the areas shown in the higher-magnification images. **(F)** Effect of genetic depletion of p53 in heat-stressed retinas. Heat-stressed retinas injected with control (top) or p53 morpholinos (bottom) 24 hours after heat stress (48 hpf). DAPI (grey) labels nuclei. Active caspase 3 labels apoptotic cells; colour represents Z depth. White dashed lines outline the embryos **(B)** or the retinas **(D, F)**. Lookup tables show minimum and maximum Z depths **(D, F)** or minimum and maximum signal values **(E)**. Scale bars, 150 μm **(B, C)**, 50 μm **(D-F)**, 10 μm for magnified regions **(E)**.

To test the previous notion that cell death following heat stress was caused by the induction of genomic instability and replicative stress, that consequently triggered p53-dependent apoptosis^46,50–52^, we performed immunostaining against ɣh2ax, a marker for DNA double-strand breaks. We observed high ɣh2ax signal in heat-stressed retinas compared to controls at 28 hpf (4 h after heat stress) (Figure 1E). At 48 hpf (24 h after heat stress), however, ɣh2ax staining was largely absent (Figure 1E). As this was the time when active caspase 3 staining was prominent, it suggested that retinal progenitors experience high replicative stress prior to the onset of apoptosis. Genetic depletion of p53 using an established morpholino approach^53^, largely rescued cell death in heat-stressed retinas at 48 hpf (24 h after heat-stress) (Figure 1F), confirming that apoptosis upon heat stress was activated primarily through the p53-pathway.

To examine the temporal progression of cell death in the retina, we characterized apoptosis progression in the 48 hours following heat stress. Apoptotic cells started to appear 4 h after heat stress (Figure S1C) and became more prominent between 10 h and 24 h after heat stress (Figure 1D, Figure S1C). By 72 hpf (48 h after heat stress), retinas showed only low levels of remaining active caspase 3 signal comparable to age-matched controls. When qualitatively evaluating retinas using DAPI staining, we noted that at 72 hpf, heat-stressed and control retinas displayed a similar arrangement of nuclear layers, indicating that retinal development progressed in both (Figure 1D). Overall, these results show that heat stress triggers extensive and widespread cell death in the zebrafish retina between 28 hpf and 48 hpf through p53-dependent apoptosis. Cell death subsided by 72 hpf and heat-stressed retinas appeared to progress in their development.

### Neurogenesis onset is not altered in heat-stressed retinas

The time window between 28 hpf and 48 hpf in which heat stress-induced apoptosis was most prevalent coincides with the onset of retinal neurogenesis in the developing zebrafish retina (Figure 1A). Previous studies have suggested that cell loss due to chemogenetic ablation in the mouse long bone or physical cell removal in the zebrafish optic vesicle can be counteracted by delaying the onset of differentiation^54,55^. To test whether apoptosis upon heat stress had a similar effect on retinal neurogenesis onset, we imaged the *Tg(ath5:GAP-RFP)* reporter line that labels the first wave of neurogenic progenitors^56–58^. We observed that, similar to the timing observed in controls, Ath5+ cells started to emerge at 28 hpf (4 h after heat stress) in 77% (N=14) of heat-stressed embryos (Figures 2A, 2B). This timely emergence of Ath5+ neurogenic progenitors was even seen when heat-stressed retinas appeared considerably smaller than age-matched controls (Figure 2A). A general delay in neurogenesis as an adaptation to compensate for extensive cell death upon heat stress was thus considered unlikely.

**Figure 2.**
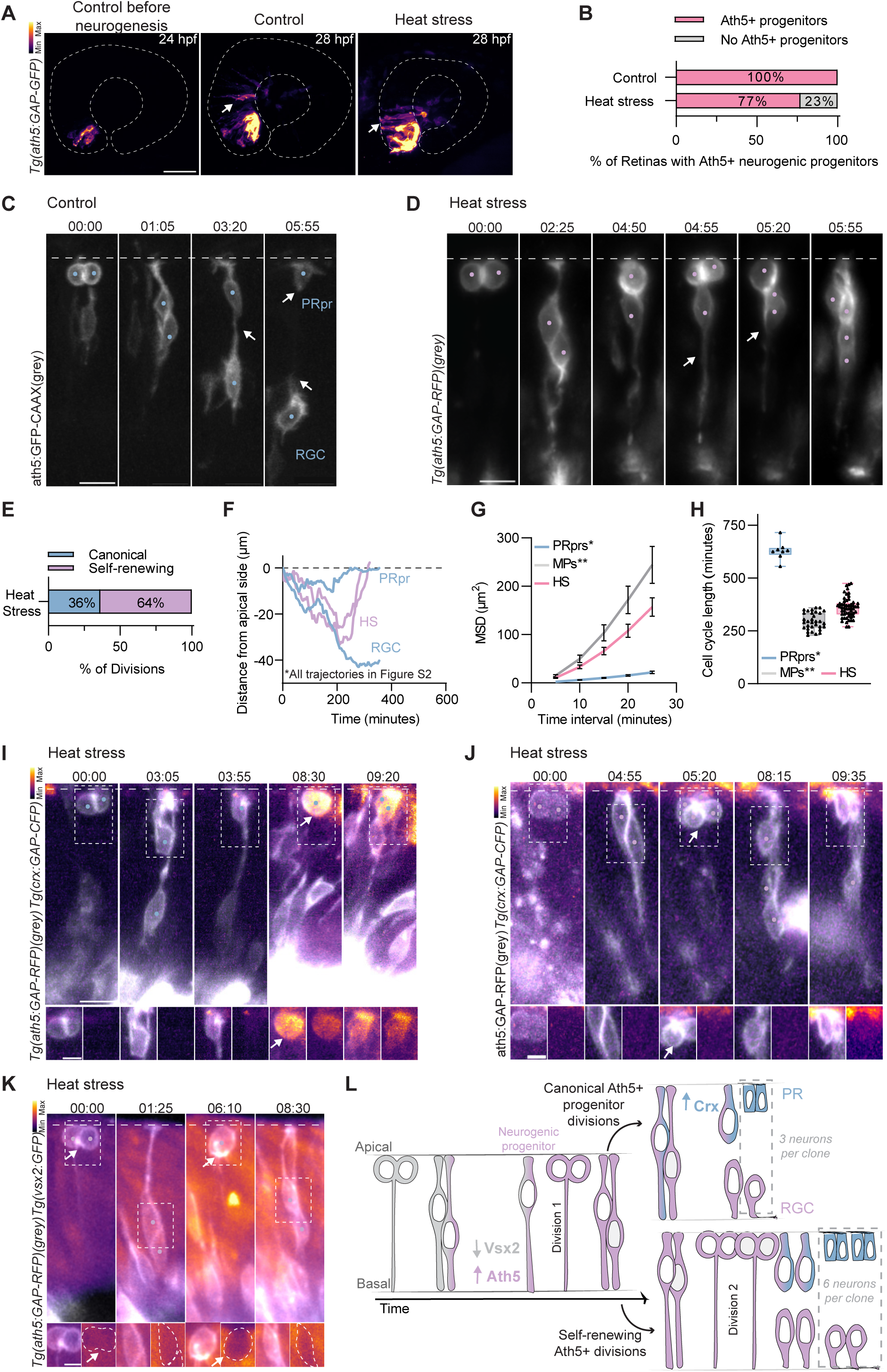
Ath5+ neurogenic progenitors undergo self-renewing divisions in heat-stressed retinas. **(A-D)** Emergence and division patterns of Ath5+ neurogenic progenitors. **(A)** Retinas at 24 hpf (before neurogenesis onset) and 28 hpf (4 h after heat stress) in control and heat-stressed embryos. Arrows indicate Ath5+ neurogenic progenitors. **(B)** Proportion of retinas containing Ath5+ neurogenic progenitors at 28 hpf in control and heat-stressed embryos (n=13 control, n=14 heat-stressed embryos). **(C)** Time series of Ath5+ neurogenic progenitor division generating a RGC (basal cell) and a PR precursor (apical cell) in a control retina. **(D)** Time series of a self-renewing Ath5+ progenitor division in a heat-stressed embryo. **(A, C-D)** *Tg(ath5:GAP-GFP)*, *Tg(ath5:GAP-RFP)* and ath5:GFP-CAAX label neurogenic progenitors. **(C-D)** Arrows indicate basal and apical processes. **(E)** Proportion of canonical and self-renewing divisions among tracked Ath5+ progenitors in heat-stressed retinas (n=72 cells, N=14 embryos). **(F)** Example trajectories of Ath5+ progenitors from birth to division in control retinas, showing the characteristic basal migration patterns of photoreceptor precursors (PRpr, blue), retinal ganglion cells (RGC, blue); and heat-stressed Ath5+ progenitors undergoing self-renewing divisions (HS, lilac). Distance from the apical side is plotted over time. All individual trajectories are shown in Figure S2A. **(G)** Mean square displacement (MSD) of the apical migration of heat-stressed Ath5+ progenitors undergoing self-renewing divisions (HS, pink, n=56 cells, α=1,678), photoreceptor precursors (PRpr, blue, n=49 cells, α=1,349) and multipotent progenitors (MP, grey n=31 cells, α=1,739) in control retinas, shown as the mean of all tracks ± s.e.m. PRpr* trajectories from *Rocha-Martins et al., 2023* and multipotent progenitor** trajectories from *Nerli et al., 2020* were re-analysed from birth to division or final position, respectively. **(H)** Cell cycle length of multipotent progenitors** (grey) and PRpr* (blue) re-analysed from Nerli et al., 2020 and *Nerli et al., 2023,* respectively, and of self-renewing divisions in heat-stressed retinas (pink). Data are presented as box plots with superimposed individual measurements (dots). Lower and upper box hinges, 25th and 75th percentiles, respectively; middle line, median; whiskers, minimum and maximum. (I-J) Time series showing expression of photoreceptor differentiation reporter in Ath5+ neurogenic progenitors. **(I)** Canonical division in a control embryo. **(J)** Self-renewing division in a heat-stressed embryo. **(I)** *Tg(ath5:GAP-RFP)* and **(J)** ath5:GAP-RFP labels neurogenic progenitors and **(I-J)** *Tg(crx:GAP-CFP)* labels PRprs. Dashed boxes indicate magnified regions shown below. In each timepoint, the left and right panels show merged RFP/CFP channels and the CFP channel, respectively. **(K)** Time series showing absence of multipotent progenitor reporter expression during self-renewing Ath5+ progenitor division in a heat-stressed embryo. *Tg(ath5:GAP-RFP)* labels neurogenic progenitors and *Tg(vsx2:GFP)* labels multipotent progenitors. Dashed boxes indicate magnified regions shown below. In each timepoint, the left and right panels show merged RFP/ GFP channels and the GFP channel, respectively. **(L)** Schematic of canonical and self-renewing divisions of Ath5+ progenitors. White dashed lines indicate the outline of the retinas **(A)** or the apical side **(C-D, I-K)**. Lookup tables show minimum and maximum Z depths **(A)** or minimum and maximum signal values **(I-K)**. **(C-D, I-K)** Time is displayed as hours:minutes. Scale bars, 50 μm **(A)**, 10 μm **(C, D, I-K)**, 5 μm for magnified regions **(I-K)**.

### Ath5+ neurogenic progenitors undergo self-renewing divisions in heat-stressed retinas

To understand how neurogenesis progresses following heat stress-induced apoptosis, we monitored the division behaviour of Ath5+ neurogenic progenitors using the *Tg(ath5:GAP-RFP)* reporter line from 28 hpf (4 h after heat stress) for up to 48 hours. We used light-sheet microscopy to minimize further stress due to phototoxicity^59^. In control embryos, Ath5+ progenitor soma moved to the apical surface and divided once, giving rise to one photoreceptor precursor and either a retinal ganglion cell (RGC) or an interneuron (Figure 2C, 2F; Video S1), as described previously^44,57,60^. Following an initial basal translocation, the photoreceptor precursor moves back to the apical surface while the sister cell moves further basally to its final laminar position, as described previously^44,60^ (Figure 2C; Video S1). This division behaviour changed in heat-stressed retinas, where we observed additional divisions in a large subset of Ath5+ progenitor daughter cells: In 64% (n=72 cells, N=14 embryos; Figure 2E) of cases, Ath5+ progenitor divisions were followed by one daughter cell (35%) or both daughters (29%) returning to the apical side and immediately dividing again (Figure 2D; Video S1; Figure S2B). Between divisions, these cells retained bipolar morphology, similar to progenitor cells (Figure 2D), a behaviour not previously reported.

To determine whether this behaviour was indeed more representative to that of progenitor cells, we analysed the movement of Ath5+ soma between apical divisions (n=56 cells, N=14 embryos; Figure S2A). The acquired data was compared to previously published data on non-neurogenic progenitor and photoreceptor precursor cell behaviours, that was acquired with the same experimental setup and re-analysed here^43,60^. Daughters of Ath5+ divisions in heat-stressed retinas exhibited directed apical movement before dividing again (Figure 2F, G), as evidenced by the supralinear mean square displacement (MSD) curve (Figure 2G). This movement differed from that of photoreceptor precursors, the only cells that canonically arise from Ath5+ divisions and return apically, which showed slower, more saltatory, translocation (Figure 2F, 2G). The apical movement of the daughters of Ath5+ divisions in heat-stressed retinas was instead more comparable to the movements of retinal progenitors (Figure 2G). Also cell cycle length of Ath5+ cells undergoing additional rounds of division in heat-stressed retinas was more similar to progenitor cell cycles, averaging 359,7 min ± 48,49 min (Figure 2H) compared to 296,1 min ± 39,89 min in progenitors^43^. This length was approximately half the duration reported for photoreceptor precursors^61^ (629,4 min ± 44,63 min; Figure 2H). That these additional Ath5+ divisions did not immediately produce photoreceptor precursors was further shown by the absence of the photoreceptor differentiation reporter Crx+ (Tg(*crx:GAP-CFP))* in Ath5+ daughter cells (Figure 2I, 2J; Video S1).

These results raised two possibilities: 1) daughter cells of extra divisions could retain neurogenic progenitor identity or 2) daughter cells could revert to a pre-neurogenic state. To differentiate between these two possibilities, we assessed the expression of a Vsx2 reporter transgene^62^, a master regulator of multipotent progenitor state that we expected to be co-expressed in Ath5+ cells in heat-stressed embryos, in case they reverted to a pre-neurogenic state. However, we found no Ath5+ cells undergoing self-renewing divisions that showed both markers upon heat stress in a *Tg(ath5:GAP-RFP; vsx2:GFP)* transgenic line (Figure 2K, n=12 cells, N=4 embryos). Thus, Ath5+ progenitors most likely undergo additional divisions without reverting to a pre-neurogenic progenitor state.

Notably, in three out of seven instances in which Ath5+ cells could be followed beyond their second division, they underwent a third division (Figure S2D; Video S1). In the remaining four cases, the daughter cells of the second division of Ath5+ progenitors produced three neurons each, PR-PR and an RGC/Interneuron as seen for canonical Ath5+ divisions (Figure S2C; Video S1). Thus, neurogenic progenitors can undergo multiple rounds of additional divisions before ultimately giving rise to their canonical neuronal offspring. These results suggest that, following cell death induced by heat stress, Ath5+ neurogenic progenitors can bypass their canonical transition to differentiating cell states. They instead undergo additional rounds of division before neuronal commitment (Figure 2L). This previously uncharacterized capacity is hereafter referred to as “self-renewing divisions”.

### Self-renewing divisions of Ath5+ neurogenic progenitors are a general response to apoptosis

To test whether self-renewing divisions of Ath5+ progenitors were a response to cell death beyond the heat stress regime, we employed a chemical-genetic ablation method (NTR-Mtz) to induce apoptosis^63,64^. Embryos were injected with RNA encoding the NTR protein at the 8-32 cell stage and subsequently treated with 10 mM metronidazole from 16 hpf - 28 hpf (Figure 3A). At 48 hpf (20 h after end of treatment), treated retinas displayed increased levels of ɣh2ax (Figure S3A) and active caspase 3 staining compared to untreated controls (Figure 3B), confirming the induction of DNA damage and apoptosis. Live imaging of Ath5+ progenitors in NTR-Mtz treated retinas (Figure 3C; Video S2) revealed that also in this regime Ath5+ cells underwent self-renewing divisions (n=10/43 cells, N=11 embryos), albeit with lower penetrance than seen in heat-stressed embryos (Figure 3L). This difference is most likely due to overall lower levels of apoptosis induced by the NTR-Mtz treatment.

**Figure 3.**
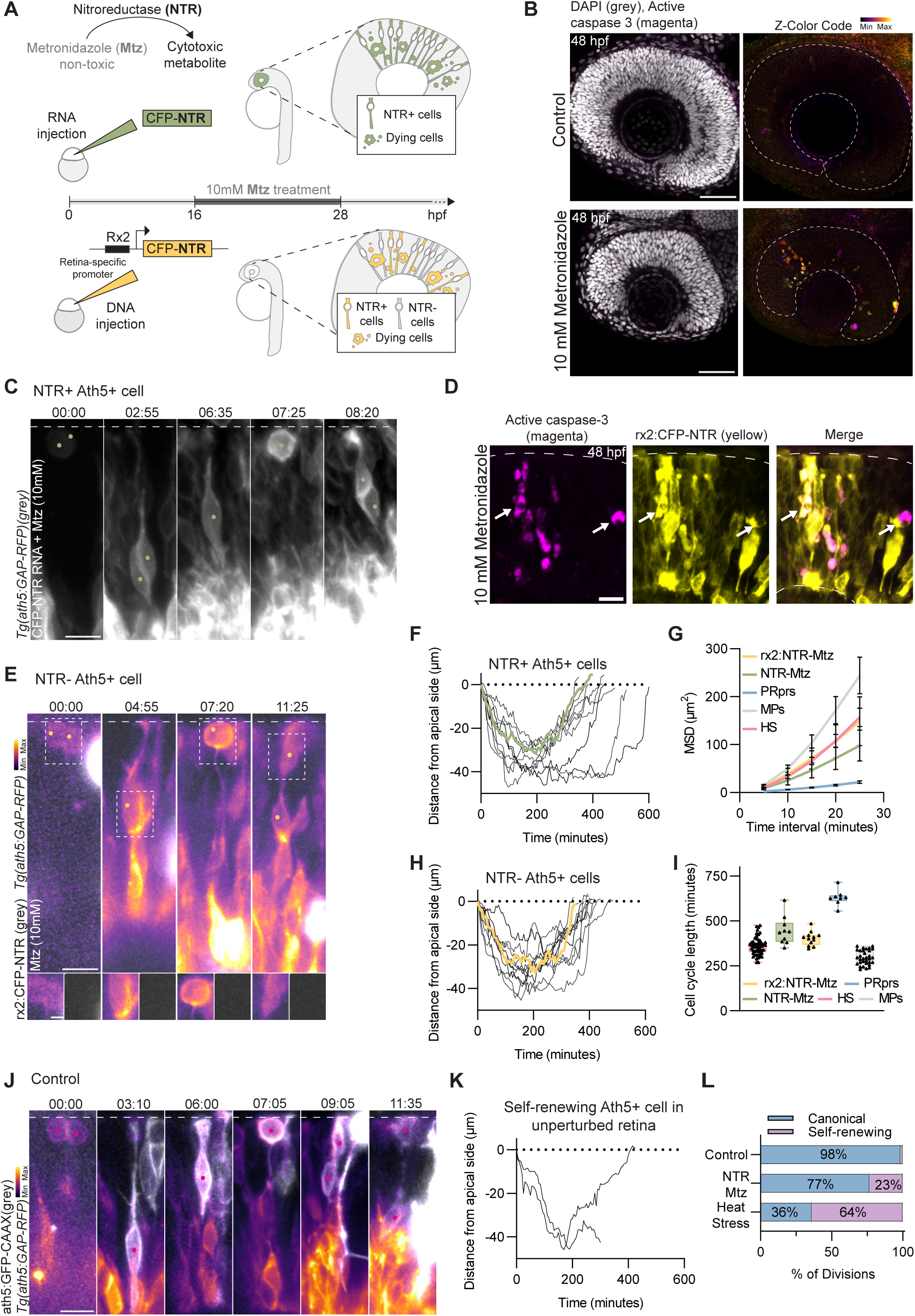
Self-renewing divisions of Ath5+ neurogenic progenitors are a non-cell-autonomous response to apoptosis. **(A)** Schematic of chemogenetic ablation using the NTR/MTZ system. Embryos injected with CFP-NTR RNA **(B-C)** or rx2:CFP-NTR DNA injections **(D-E)** were treated with 10 mM Mtz between 16 hpf and 28 hpf to induce cell death in NTR-expressing cells. **(B)** Spatial distribution of apoptotic cells in retinas at 48 hpf in CFP-NTR RNA-injected control (top) and Mtz-treated (bottom) embryos. DAPI (grey) labels nuclei. Active caspase 3 labels apoptotic cells; colour represents Z depth. **(C)** Time series of a self-renewing Ath5+ progenitor in a CFP-NTR RNA-injected/Mtz-treated embryo. *Tg(ath5:GAP-RFP)* (grey) labels neurogenic progenitors. **(D)** Spatial distribution of apoptotic cells in retinas at 48 hpf in rx2:CFP-NTR DNA-injected/Mtz-treated embryos. Active caspase 3 (magenta) labels apoptotic cells; CFP-NTR-expressing cells (yellow). Arrows indicate apoptotic CFP-NTR positive cells. **(E)** Time series of a self-renewing Ath5+ NTR-progenitor in a rx2:CFP-NTR DNA-injected/Mtz-treated embryo. *Tg(ath5:GAP-RFP)* labels neurogenic progenitors. Dashed boxes indicate magnified regions shown below. In each timepoint, the left and right panels show RFP and CFP channels, respectively. **(F)** Trajectories of self-renewing Ath5+ progenitors in CFP-NTR RNA-injected/Mtz-treated embryos. A representative trajectory is highlighted in green (n=10 cells, N=10 embryos). Distance from the apical side is plotted over time. **(G)** Mean square displacement (MSD) curves of apical migration of self-renewing Ath5+ progenitors in CFP-NTR RNA-injected/Mtz-treated embryos (NTR-Mtz, green, n=10 cells, α=1,515) and heat-stressed embryos (HS, pink, n=56 cells, α=1,678), Ath5+ NTR-progenitors in rx2:CFP-NTR DNA-injected/Mtz-treated embryos (yellow, n=12 cells, α=1,565), photoreceptor precursors (PRpr, blue, n=49 cells, α=1,349) and multipotent progenitors (MP, grey n=31 cells, α=1,739) in control embryos, shown as the mean of all tracks ± s.e.m. **(H)** Trajectories of self-renewing Ath5+ NTR- progenitors in a rx2:CFP-NTR DNA-injected/Mtz-treated embryo. Distance from the apical side is plotted over time. A representative trajectory is highlighted in yellow. (n=12 cells, N=3 embryos). **(I)** Cell cycle length of self-renewing Ath5+ progenitors in CFP-NTR RNA-injected/Mtz-treated (green) and heat-stressed embryos (pink), Ath5+ NTR-progenitors in a rx2:CFP-NTR DNA-injected/Mtz-treated embryo (yellow), multipotent progenitors (grey) and PRpr (blue) in control embryos. Data are presented as box plots with superimposed individual measurements (dots). Lower and upper box hinges, 25th and 75th percentiles, respectively; middle line, median; whiskers, minimum and maximum. **(J)** Time series of a self-renewing Ath5+ division observed in an unperturbed retina. ath5:GFP-CAAX (grey) and *Tg(ath5:GAP-RFP)* label neurogenic progenitors. **(K)** Trajectory of the self-renewing Ath5+ progenitor shown in panel J. Distance from the apical side is plotted over time. **(L)** Proportion of canonical versus self-renewing divisions among tracked Ath5+ progenitors in control (n=59 cells, N=14 embryos), NTR-Mtz treated (n=43 cells, N=11 embryos), and heat-stressed retinas (n=72 cells, N=14 embryos). MSDs **(G)** and cell cycle **(I)** lengths of PRpr, multipotent progenitors and self-renewing Ath5+ progenitors in heat-stressed embryos replotted from Figure 2 for comparison. White dashed lines indicate the outline of the retina **(B, D)** or the apical side **(C, E, J)**. Lookup tables show minimum and maximum Z depths **(B)** or minimum and maximum signal values **(C, E, J)**. **(C, E, J)** Time is displayed as hours:minutes. Scale bars, 50 μm **(B)**, 20 μm **(D)**, 10 μm **(C, E, and J)** and 2 μm for magnified regions **(E)**.

As seen in embryos exposed to heat stress, self-renewing Ath5+ cells displayed a directed apical movement before mitosis, as shown by the supralinear slope of their MSD curve (Figure 3F, 3G). Further, their average cell cycle length (445 min ± 78 min) was shorter than that reported for photoreceptor precursors^61^ (629,4 min ± 44,63 min; Figure 3I).

These findings show that in different apoptosis-inducing regimes, neurogenic Ath5+ progenitors undergo self-renewing divisions, indicating that this change in behaviour is a general response to cell death.

### Self-renewal of Ath5+ progenitors is a non-cell-autonomous response to apoptosis

To understand if cells that are not directly affected by NTR-induced apoptosis undergo self-renewing divisions, we generated a CFP-NTR DNA construct under the control of the retina-specific promoter Rx2. Mosaic expression of NTR was achieved by one-cell-stage injection of the DNA construct. Cell death was induced by treatment of embryos with 10 mM metronidazole from 16 hpf - 28 hpf as described above (Figure 3A). We found that all caspase 3-positive cells co-express CFP-NTR (Figure 3D), confirming induction of apoptosis exclusively in Rx2 expressing cells. When following ath5:GAP-RFP positive, rx2:CFP-NTR negative progenitors (Figure 3E; Video S2), we observed Ath5+ cells that were not directly affected by NTR and underwent self-renewing divisions (n=12 cells, N=3 embryos; Figure 3E). MSD analysis confirmed that these cells displayed a directed apical movement prior to mitosis as noted in the other apoptosis-inducing regimes (Figure 3H, 3G). Like self-renewing Ath5+ progenitors in heat stress and NTR-Mtz treated embryos, their average cell cycle length (401,7 min ± 39,16 min; Figure 3I) was shorter than that reported for photoreceptor precursors^61^ (629,4 min ± 44,63 min; Figure 3I).

The finding that unperturbed Ath5+ progenitors can undergo self-renewing divisions prompted us to ask whether such divisions could also be found in rare occasions in embryos that were not exposed to elevated stress levels. To address this question, we tracked Ath5+ neurogenic progenitor divisions in non-perturbed retinas (n=59 cells from N=14 embryos). We found one Ath5+ neurogenic progenitor cell undergoing a self-renewing division (Figure 3J, 3L; Video S2). The translocation trajectory of this cell was similar to what was seen for the self-renewing divisions in heat-stressed, NTR-Mtz and rx2:NTR-Mtz treated retinas (Figure 3K). Taken together, these results suggest that self-renewing capacity is an intrinsic property of Ath5+ neurogenic progenitors, which occurs sporadically in control situations but is upregulated in response to induced apoptosis.

### The embryonic zebrafish retina recovers from extensive apoptosis and acquires visual function

The fact that a large portion of Ath5+ progenitors undergo additional rounds of division upon cell loss means that these cells effectively produce larger clones (Figure 2L). Based on the frequency of self-renewing divisions observed in heat-stressed retinas, we estimated that each Ath5+ neurogenic progenitor produces on average 4.2 neurons instead of 3 in the canonical scenario. This corresponds to an approximately 40% increase in clonal output. Thus, we asked whether this increased cell production could enable the restoration of tissue formation and visual function. This notion was consistent with the finding that heat-stressed retinas seemed to progress in development despite extensive cell loss (Figure 1D).

We followed retinal size in heat-stressed versus control embryos measuring the areas of eyes between 8 hours after heat stress (32 hpf) until 5 dpf (Figure S4A). Over this period, the eye area of heat-stressed retinas increased by 135% (compared to 111% in controls), showing that growth continues despite significant cell loss (Figure S4A). To understand 3D growth dynamics of control versus heat-stressed retinas, we reconstructed retinal volumes in IMARIS software using tagged laminin to outline the retinal surface of live embryos at 48 hpf and 72 hpf (24 h and 48 h after heat-stress). This analysis showed that the growth rate was similar between heat-stressed retinas (n=16; 60385 μm^3^/hour ± 29944 μm^3^/hour) and controls (n=15; 56389 μm^3^/hour ± 18462 μm^3^/hour; Figure 4B). Analysis of fixed embryos confirmed this notion. Heat-stressed retinas were on average 50% smaller than controls at 48 hpf (24 h after heat stress; Figure 4A, 4C). Retinal volume increased 2.19x over the subsequent 3 days, compared to 1.75x in controls (4d after heat stress; Figure 4C). At 5 dpf, heat-stressed retinas reached on average 64% of control volume, with some showing volumes equal to controls. Thus, despite extensive cell death, heat-stressed retinas continue to grow at comparable or even slightly higher rates than controls.

**Figure 4.**
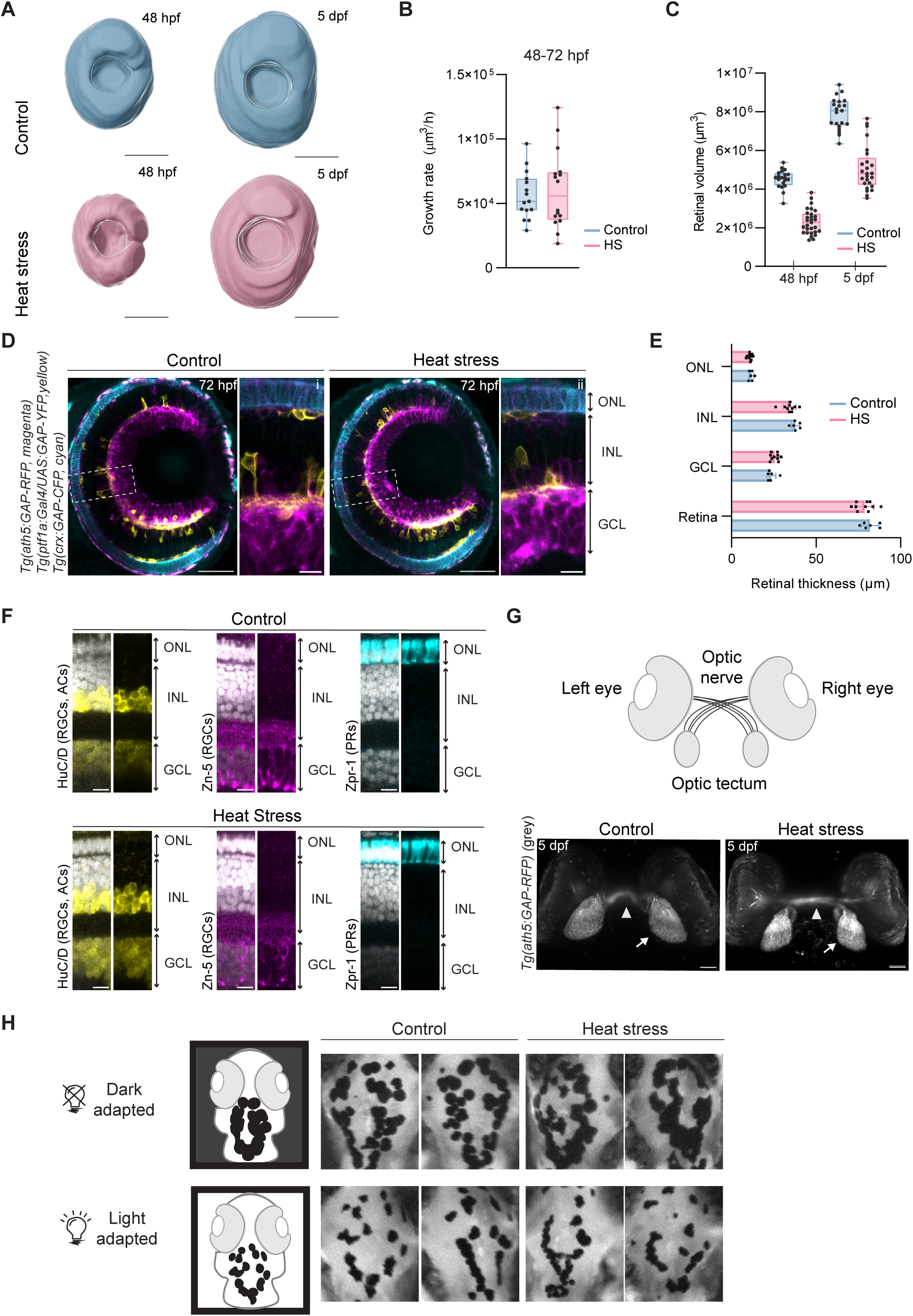
Heat-stressed retinas grow, recover tissue architecture and visual function. **(A)** Three-dimensional surface reconstructions of control (blue; top) and heat-stressed (pink; bottom) retinas at 48 hpf (left) and 5 dpf (right). **(B)** Growth rate of retinal volume between 48 hpf and 72 hpf in control (blue, n=15) and heat-stressed embryos (pink, n=16). P=0.9224 for comparison between control and heat-stress embryos using Mann–Whitney test (two-sided). **(C)** Retinal volume in fixed control (blue, n=20, n=21) and heat stress (pink, n=29, n=25) 1 d and 4 d after heat stress. P<0.0001 for comparison between control and heat stress embryos at each timepoint using Mann–Whitney test (two-sided). **(B-C)** Data are presented as box plots with superimposed individual measurements (dots). Lower and upper box hinges, 25th and 75th percentiles, respectively; middle line, median; whiskers, minimum and maximum. **(D)** Neuronal lamination in control (left) and heat-stressed (right) retinas at 72 hpf. *Tg(ath5:GAP-RFP)* labels RGCs (magenta), *Tg(ptf1a:Gal4/UAS:GAP-YFP)* labels inhibitory neurons (yellow) and *Tg(crx:GAP-CFP)* labels photoreceptors (cyan). Dashed white boxes outline magnified regions. **(E)** Thickness of the retina and individual layers (ganglion cell layer (GCL), inner nuclear layer (INL), and outer nuclear layer (ONL) in control and heat-stressed retinas at 72 hpf. Mean and SD are indicated. Individual data points are shown (n=6 control, n=10 heat-stressed embryos). P-values for comparison between control and heat-stressed embryos using Mann–Whitney test (two-sided) are found in Table 1. **(F)** Neuronal maturation in control (top) and heat-stressed (bottom) retinas at 5 dpf. Zn-5 labels retinal ganglion cells (RGCs, magenta), HuC/D labels RGCs and amacrine cells (ACs, yellow), Zpr-1 labels photoreceptors (PRs, cyan) and DAPI (grey) labels nuclei. **(G)** Schematic and representative images of the optic nerves and neuronal projections to the tectum labelled by *Tg(ath5:GAP-RFP)* (grey) in control (left) and heat-stressed (right) embryos at 5 dpf. **(H)** Visual background adaptation assay in control (left) and heat-stressed (right) larvae at 5 dpf. Schematics show the expected pigmentation in dark (top) and light (bottom) conditions. Brightfield images of individual heat-stressed and control larvae. Scale bars, 100 μm **(A)**, 50 μm **(D, G)**, 10 μm **(F)**.

**Table 1.** Layer thickness measurements for control and heat stress conditions.

|  | **Total thickness (μm)** | **ONL** | **INL** | **GCL** | **N** |
| --- | --- | --- | --- | --- | --- |
| Control | 81,91 ± 5,04 | 11,76 ± 1,37 | 37,85 ± 2,46 | 23,56 ± 2,65 | 6 |
| Heat stress | 78,72 ± 5,42<br>(P=0,3531) | 10,91±1,22<br>(P=0,3529) | 34,31 ± 3,76<br>(P=0,0447) | 25,73 ± 2,31<br>(P=0,1179) | 10 |

Mean and SD are shown for each measurement. P-values of comparison with control using Mann–Whitney test (two-sided) are shown for each measurement. N refers to the number of embryos analysed.

At peak apoptosis stages (24 h after heat stress), DAPI staining revealed severe retinal disorganization and misoriented nuclei (Figure 1D). By 72 hpf, however, heat-stressed retinas seemed laminated, indicating that restoration of retinal architecture had occurred (Figure 1D). When analysing cell type distribution in the Spectrum of fates transgenic line^65^ that allows us to unambiguously differentiate all major retinal neuronal types, we noted that heat-stressed retinas at 72 hpf contained all neuronal types, arranged in their correct layers (Figure 4D), which showed similar thickness to control embryos (Figure 4E, Table 1). Further, at 5 dpf, markers of maturing photoreceptors (Zpr-1), retinal ganglion cells (Zn-5) and basal neurons (HuC/D) were seen in their appropriate layers (Figure 4F). Thus, compensation to cell loss appears to support neuronal differentiation and the acquisition of correct tissue architecture. To understand if this restoration of tissue architecture went beyond the retina and led to the generation of visual function, we probed whether heat-stressed retinas generate an optic nerve. 3D reconstruction of *Tg(ath5:GAP-RFP)* embryos showed that 88% of heat-stressed retinas (n=30 control, n=32 heat-stressed embryos) featured an optic nerve at 5 dpf (4 days after heat stress; Figure 4G, Figure S4B) morphologically similar to what was seen in controls. To assess visual function, we used the background adaptation response, in which zebrafish adjust their skin pigmentation to ambient brightness in a vision-dependent manner^66,67^. At 5 dpf, heat-stressed larvae adapted melanocyte morphology to light and dark environments indistinguishably from controls (Figure 4H), demonstrating that these fish can indeed perceive visual information.

Thus, heat-stressed retinas that suffered massive cell death continue to grow and, by larval stage, contain all neuronal types in the correct layers. This robustness to extensive cell loss allowed to enable light perception, suggesting that the compensatory response can support visual function acquisition after severe developmental perturbation.

## Discussion

This study investigated aspects of cellular plasticity that underlie robustness against cell loss in the developing CNS. The rapid development and transparency of the zebrafish retina allowed us to use high-resolution *in vivo* imaging to monitor the response of cells and tissue to induced apoptosis from onset of damage through morphogenesis. We find that upon heat stress or chemogenetic ablation that induced widespread cell death, a large fraction of the Ath5+ neurogenic progenitor population undergoes additional rounds of division instead of immediate neuron production. This shift in neurogenic progenitor behaviour is a non-cell autonomous response to apoptosis that occurs concurrently with ongoing retinal growth and neuronal differentiation. These self-amplifying Ath5+ divisions increase clonal output in retinas that experienced widespread cell loss. This provides a substantial source of replacement cells that help to preserve retinal morphogenesis and ultimately visual function.

The finding that Ath5+ neurogenic progenitors have the capacity to undergo self-renewing divisions extends prior knowledge of neural progenitor behaviour in the developing vertebrate retina. Previous studies in mice and zebrafish have shown that equipotent retinal progenitors generate highly variable clonal outputs through stochastic fate decisions, yet reliably produce retinas of consistent size, composition and function^39,40,68^. This suggested that the developing retina can buffer single cell variability and thereby ensure predictable outcomes at the tissue level. Recent work in zebrafish further revealed unexpected plasticity in these fate decisions. When key fate determinants were removed, early neurogenic progenitors did not arrest at this progenitor stage or undergo cell death but instead widened their spectrum of fate possibilities and even generated neuronal lineages that do not occur in control conditions^44^. With this study we further extend the known plasticity of early neurogenic progenitors by showing that, when the developing retina experiences excessive cell loss, Ath5+ progenitors bypass their canonical transition to a differentiating cell state and instead undergo additional rounds of division. This produced an estimated ∼40% increase in clone size that likely aids to buffer retinogenesis against cell loss. So far, it is unclear whether this adaptive plasticity is restricted to Ath5+ progenitors or whether other progenitor states such as multipotent proliferative progenitors and committed precursors can also adjust their proliferative behaviour. Additional buffering mechanisms against cell loss are likely, as earlier work in zebrafish in which cells were physically removed at the optic vesicle stage showed a delayed onset of neurogenesis^55^. This suggests that cell loss at earlier developmental stages can change the division behaviour of multipotent progenitors. Together, these findings raise the possibility that the developing zebrafish retina uses distinct pools of progenitor plasticity to compensate for cell loss at different stages of development. Additional studies are necessary to explore how adaptive plasticity of different progenitor states collectively supports robust tissue growth.

We show that mosaic apoptosis is sufficient to trigger self-renewal of unperturbed Ath5+ progenitors. Self-renewing divisions are transient, with progenitors subsequently resuming neurogenesis and producing the same cell types as seen in controls. This suggests that regulatory mechanisms exist that activate self-amplification in response to cell death, but, importantly, also shut it down once the tissue recovered. Future research will need to explore how these progenitors sense cell death in the surrounding tissue and how signals are transduced into transient altered proliferative behaviour. It will also be important to unveil the nature of such signals. Some mechanisms have been suggested by studies of compensatory growth in developing tissues and during adult regeneration: in the *Drosophila* imaginal wing and eye discs, apoptotic cells stimulate proliferation of surviving neighbours through secretion of mitogens^23,24^. In the adult retina of zebrafish, dying neurons release tumour necrosis factor alpha (TNFα), which stimulates reprogramming of quiescent stem cells, a crucial step in regeneration^69,70^. MDCK epithelial cells that undergo apoptosis in vitro can generate forces that trigger compensatory proliferation in neighbouring cells via YAP-mediated mechanotransduction^71^. Defining whether these and/or other pathways are involved in the adaptive plasticity of Ath5+ progenitors will be essential to understand how their proliferative potential is restrained during normal development to prevent overgrowth yet remains tuneable under stress. It will be equally important to understand the mechanisms that remove dying cells, as caspase positive cells are not observed at later developmental stages. Possible mechanisms include apoptotic cell removal by professional phagocytes^72,73^ or neuroepithelial cells^74^. The pathways operating in this context remain to be determined.

Self-renewal of Ath5+ progenitors was also observed in one unperturbed embryo. This suggests that this behaviour is not exclusive to the apoptosis-induced state. This notion is consistent with previous large-scale genetic lineage tracing studies of Ath5+ progenitors in unperturbed embryos, which identified rare large clones composed of 5 cells or more (0.1% of 484 cells)^75^. This would be beyond the output of the canonical behaviour of Ath5+ progenitors because each neurogenic division typically only produces one photoreceptor precursor that gives rise to two photoreceptors (PR-PR) and either a post-mitotic neuron (RGC/AC) or a horizontal cell precursor that gives rise to two horizontal cells (HC-HC), thus never producing a clone size bigger than 4. These rare larger clones are instead compatible with the self-renewing behaviour observed in this study. It is therefore tempting to speculate that the plasticity seen here for Ath5+ progenitors reflects a form of cryptic cellular variability. This could be analogous to cryptic genetic variation, a well-established concept in evolutionary biology that is used to describe hidden genetic potential which only becomes relevant under stress conditions such as ecological perturbations and disease^76–78^. By analogy, self-renewal of Ath5+ progenitors would represent a latent cellular potential that can be activated upon developmental perturbations. Our discovery of this unexpected behaviour in one progenitor population that has been widely studied for decades^43,44,57,60,79,80^ argues that progenitor variability in the developing CNS is far from being understood. We hypothesize that similar hidden potentials may exist in other regions of the developing CNS, waiting to be uncovered. In the mouse brain, lineage-tracing studies have shown that some cell state transitions as well as fate decisions are stochastic^41^, raising the possibility that progenitors in other systems also retain a degree of plasticity that can be modulated under stress. Consistent with this idea, previous work revealed that induction of cell death in the embryonic cerebral cortex of mice resulted in increased proliferation of intermediate progenitors that normally undergo self-consuming divisions^22^. Together with the findings presented here, this points towards the possibility that latent plasticity in intermediate progenitor states may be prevalent more broadly across the CNS to buffer against cell loss during tissue differentiation. Thus, studying different areas of the CNS upon developmental perturbations will be important to determine whether and how hidden cellular variability contributes to the robustness of organ development.

Self-renewal of Ath5+ progenitors is particularly pronounced after acute heat stress, a relevant physical challenge for externally developing embryos. We find that even when such heat stress generates high level of cell death, retinal development and acquisition of visual function proceed. We speculate that plasticity phenomena analogous to what is observed for Ath5+ progenitors could confer resilience in disease contexts associated with unwanted cell death. In line with this idea, it has been hypothesized that compensatory proliferation, and more broadly, the regenerative capacity of developing tissues, might contribute to the incomplete penetrance observed in *Drosophila* models of microcephaly and microphthalmia^33^. Similarly, in X-linked disorders in humans associated with microphthalmia and retinal defects, skewed X-chromosome inactivation has been proposed to reflect selective elimination of cells expressing the mutant allele and compensatory expansion of unaffected cells, potentially explaining why some mutation carriers show little or no tissue defect^34^. So far, the cellular sources of this compensatory expansion remain unknown. Our findings point to progenitor plasticity as a potential mechanism of buffering against cell loss. Testing this idea will require comparative studies in disease models such as human retinal and brain organoids that are accessible for live imaging to elucidate whether adaptive plasticity of neural progenitors can mitigate developmental defects in pathological contexts.

## Material and Methods

### Zebrafish husbandry

Wild-type zebrafish and transgenic lines (*Danio rerio*; AB, TL and TU strains) were maintained at 28.5 °C (pH 7.0, conductivity 1000 μS/cm, 14 h light – 10 h dark cycle) in a recirculation life support system (Tecniplast). Embryos were raised at 28.5 °C or 32 °C in E3 medium with 0,00002% methylene blue (MB). Staging was performed in hours post-fertilization (hpf) according to Kimmel et al., 1995. Medium was changed daily and supplemented with 0.2 mM 1-phenyl-2-thiourea (PTU) (Acros Organics) from 6 hpf to prevent pigmentation. Zebrafish embryos for experiments were used from 24 to 120 hpf. All animal work was conducted in accordance with institutional standard operating procedures, under the licensing of the DGAV (Direção Geral de Alimentação e Veterinária, Portugal) in accordance with the Portuguese Decree Law n°113/2013, as well as the European Union Directive 2010/63/EU and the German Animal Welfare Act.

### Zebrafish transgenic lines

For retinal volume reconstructions, laminin reporter lines *Tg(lama1:lama1-sfGFP)* and *Tg(lama1:lama1-mKate2)* were used. To visualize emerging neurogenic progenitors, Ath5+ neurons and PRs, *Tg(ath5:GAP-RFP), Tg(ath5:GAP-GFP)* and double transgenic *Tg(ath5:GAP-RFP, crx:GAP-CFP)* lines were used. To visualize all retinal neurons or retinal progenitors, the triple transgenic line *Tg(crx:GAP-CFP)*, *Tg(ath5:GAP-RFP)* and *Tg(ptf1a:Gal4/UAS:GAP-YFP)* or *Tg(vsx2:GFP)* line were used, respectively.

### DNA constructs and cloning

The rx2:CFP-NTR expression plasmid was assembled using Gateway cloning (Thermo Fisher Scientific) with the Tol2 kit ^82^. The pCS2 vector containing the CFP-tagged NTR sequence was used for generating the pENTR(L1-L2)-CFP-NTR by PCR using Phusion polymerase (New England Biolabs) and primers with an ATT recombination site (shown in lower case): forward 5′-ggggacaagtttgtacaaaaaagcaggctggGAATTCCGGTCGCCACCATG-3’ and reverse 5’-ggggaccactttgtacaagaaagctgggtcTAGCCGGGCAGATGCCCGGC-3’. This construct was then combined with p5ENTR(L4-R1)-Rx2^83^ and pDEST(R4-R2)-Tol2pA^84^ to generate the Tol2-Rx2:CFP-NTR sequence. The ath5:GFP-CAAX^79^ was used to label Ath5+ neurogenic progenitors.

### DNA, RNA and morpholino injections

To mosaically label cells in the zebrafish retina, one-cell stage embryos were injected with purified plasmid DNA diluted in ddH_2_O supplemented with 0,05% phenol red (Sigma Aldrich). Injection volumes of the 15 ng/μL-30 ng/μL constructs ranged from 0.5-1 nl. mRNA was synthesized using the Thermo Fisher Scientific mMESSAGE mMACHINE SP6 Transcription Kit. To knockdown p53, the morpholino targeting p53 (Gene Tools, 5’-GCGCCATTGCTTTGCAAGAATTG-3’) was injected into the yolk at 4 ng per embryo with co-injection of GFP-Ras RNA at 50 pg per embryo.

### Embryo survival analysis

Control and heat-stressed embryos were maintained in separate Petri dishes containing 20-30 embryos each. Embryos were visually inspected every 24 h following treatment and dead embryos were manually counted and removed from the plates.

### Cell death induction by heat stress

To induce cell death in the developing zebrafish CNS, we subjected 24 hpf embryos to 42 °C for 45 minutes. Plastic Petri dishes containing 30-60 embryos with 40 ml of E3 supplemented with methylene blue and sealed with parafilm were placed in a pre-heated water bath. Petri dishes were allowed to cool down for 10-15 minutes at room temperature. Next, embryo medium was exchanged, PTU was added and embryos were raised at 28 °C.

### Cell death induction by NTR/Metronidazole system

For global chemogenetic cell ablation, embryos at 8–32-cell-stage were injected with 0.5-1 nL of 250 ng/μL of CFP-NTR RNA and grown at 32 °C. For retina-specific chemogenetic cell ablation, one-cell-stage embryos were injected with 20 ng/μL of rx2:CFP-NTR DNA, together with 50 ng/μL of Tol2 transposase RNA, as it improved expression efficiency.

At 16 hpf, embryos were dechorionated and treated with a 10 mM metronidazole (Sigma Aldrich) solution prepared in E3 supplemented with methylene blue solution with 2% DMSO for 12 h at 28 °C. To remove metronidazole, embryo medium was replaced three times, PTU was added and embryos were raised at 28 °C.

### Whole-mount staining of zebrafish embryos

Zebrafish embryos were dechorionated manually and fixed in 4% paraformaldehyde (PFA) in Phosphate-Buffered Saline (PBS) overnight at 4 °C or at room temperature for 2-3 h shaking at 100rpm. Embryos up to 48 hpf were washed three times for 10 minutes with 0.5% Triton X-100 in PBS (PBS-T 0.5%). For permeabilization, they were first washed with PBS, followed by ddH_2_O, before incubating with ice-cold acetone for 10-15 min at −20 °C. Acetone was rinsed twice with ddH_2_O, once with PBS, followed by three washes for 10 min with PBS-T 0.5%. Embryos older than 48 hpf were washed four times for 15 minutes with 0.8% PBS-T. For permeabilization, they were incubated with 0,25% Trypsin-EDTA in PBS on ice for different periods of time depending on the developmental stage (17 min for 72 hpf and 20 min for 96 and 120 hpf). The permeabilization solution was replaced with PBS-T 0.8% and embryos were incubated for 30 min on ice before rinsing twice with PBS-T 0.8%. For all developmental stages, blocking was performed with 10% donkey serum in PBS-T 0.8% for 2-3 h at room temperature with slight agitation, or overnight at 4 °C. Primary and secondary antibodies were diluted in 1% donkey serum in PBS-T 0.8%. Embryos were incubated with the following primary antibodies for 72h at 4°C: active caspase-3 (BD Biosciences or BD Pharmingen) 1:200, Histone H2A.XS139ph (phospho Ser139, GeneTex) 1:200, Zpr-1 (ZIRC) 1:200; Zn-5 (ZIRC) 1:50; anti-GFP (Proteintech) 1:100; anti-HuC/HuD (Invitrogen) 1:250. Embryos were washed five times for 30 min with PBS-T 0.8% and incubated for 48 h with the appropriate fluorescently labelled secondary antibody (Molecular Probes) at 1:500 and DAPI 1:1,000 (1 µg/mL) (Thermo Fisher Scientific). Finally, embryos were washed four times for 15 min with PBS-T 0.8% and stored in PBS at 4 °C until imaging.

### Image acquisition

#### Light-sheet imaging

Live imaging of Ath5+ neurogenic progenitors was performed using Light-sheet Z.1 (Carl Zeiss Microscopy) operated with ZEN 2014 SP1 software (v9.2.8.60) (black edition), as previously described^85^. Embryos were mounted in 0.6% low-melting agarose (Sigma-Aldrich) prepared in E3 medium, supplemented with 240μg/ml MS-222 (Sigma-Aldrich). The sample chamber was filled with E3 medium supplemented with 0.04% MS-222 and 30.4 μg/ml 1-phenyl-2-thiourea and maintained at 28.5 °C during the imaging session.

A single view spanning the entire volume of each eye (around 150 Z slices with a step size of 1 μm) was acquired every 5 minutes for up to 48 h using dual-sided illumination by 10x/0.2 objectives (Carl Zeiss Microscopy). Detection was performed with a Plan-Apochromat 20×/1.0 W objective (Carl Zeiss Microscopy) and two Edge 5.5 or 4.2 sCMOS cameras (PCO).

#### Laser scanning confocal microscopy

Dechorionated samples were imaged with a Zeiss LSM980 Airyscan2 inverted microscope, equipped with two PMT and one GaAsP detector using a 40×/1.1 C-Apochromat water immersion objective from Zeiss for retinal acquisitions and 10x/EC Plan-Neofluar objective for whole-embryo acquisitions. Embryos were mounted in 1% low-melting agarose in glass-bottomed dishes (35 mm; MatTek Corporation) and imaged at room temperature. The microscope was operated using the ZEN Blue v3.3 (Zeiss) software. Acquired Z-stacks encompassed the centre or the entire retinal volume (around 50 Z slices with a step size of 1-1.5 μm).

### Image processing and analysis

Image data were processed in ZEN Black and/or Fiji. Spatial drift of live-imaging data was corrected using the Manual Drift Correction plug-in (http://imagej.net/Manual_drift_correction_plugin, ImageJ). Data were visualized in 3D using Imaris (v10.0.0).

#### Cell tracking

Neurogenic progenitors were identified based on the expression of the ath5 reporter and progenitor morphology such as the presence of apical and basal attachments. In cases where sparse labelling by mosaic reporter expression facilitated tracking of individual cells, progenitor identity was confirmed by the presence of *Tg(ath5:GAP-RFP)* signal. The progeny of Ath5+ cells was followed from birth to cell division or final positioning in maximum projections of substacks of the time-lapse datasets using Fiji. Trajectories were obtained by tracing the centre of the cell using the ImageJ plug-in MtrackJ (https://imagej.net/plugins/mtrackj). Trajectories in y were offset to the cells’ initial position after birth.

#### Retinal volume reconstructions

To reconstruct the retinal surface, *Tg(lama1:lama1-sfGFP)*^86^ or *Tg(lama1:lama1-mKate2)*^86^ signal from live and fixed embryos imaged in the Light-sheet Z.1, was manually outlined (every 5-10 slices) across the entire retinal volume. Volume of the segmented surface was calculated using IMARIS software.

#### Retinal thickness measurements

Retinal measurements were performed manually using the Fiji line tool on confocal Z-stacks of controls and heat-stressed retinas at 72 hpf. The *Tg(ptf1a:Gal4; UAS:YFP; ath5:GAP-RFP; and crx:GAP-CFP)* line was used for discrimination of retinal layers. Retinal thickness was measured on three different Z-planes of the central region of the tissue, and the average measurement was plotted for each biological replicate. The outer nuclear layer (ONL) was assigned as the distance between the apical surface of the retina and the outer plexiform layer; the inner nuclear layer (INL) as the distance between the inner plexiform layer and the outer plexiform layer; and the ganglion cell layer (GCL) was assigned as the layer between the inner plexiform layer and the basal lamina, as done previously^44^.

### Visual background adaptation assay

To assess retinal perception of light we adapted the visual background adaptation (VBA) assays described in^66,67,87^. 4 dpf control and heat-stressed larvae were split into two groups: one kept on a white background overnight under light, and another kept on a black background overnight, in the dark. The VBA response was evaluated the next morning, immediately after the start of the normal light cycle. The animals used for imaging and VBA quantification were euthanised in MS-222 and immediately fixed in 4% PFA for 1 h. To assess melanophore morphology, 5 dpf larvae were washed in PBS, and imaged under a stereomicroscope.

### Quantification and statistical analysis

#### Analysis of kinetics of cellular movements

MSDs were calculated from the 25 min of movement prior to apical mitosis in Excel (Microsoft) using the open-source computer program DiPer^88^. To estimate directional persistence, the α-value was obtained from the slope of the log-log plots of MSDs and time interval. For photoreceptors and non-neurogenic multipotent progenitors, trajectories were re-analysed in the same manner from data previously published by the lab^43,60^.

#### Clone size estimation

To estimate the expansion of Ath5+ neurogenic progenitors in heat-stressed retinas, we considered the relative frequency of each mode of division observed (Figure S2B) and their expected clone size. For canonical neurogenic divisions, we considered an output of three cells: one photoreceptor precursor that gives rise to two photoreceptors (PR-PR) and a post-mitotic neuron (RGC/AC). Because horizontal cell precursors represent a minor fraction of retinal neurons^44^, their contribution was not included. Divisions in which one daughter cell self-renewed were assigned a clone size of four cells (one committed precursor and two neurons), whereas divisions in which both daughter cells self-renewed were assigned an expected output of six cells (two committed precursors and two neurons). The average expected clonal output was then calculated as a weighted sum of these clone sizes according to the observed frequency of each division mode.

#### Statistics

Statistical tests were performed using GraphPad Prism v.9.4.0. All statistical tests used are indicated in the figure or table legends and the definitions of error bars. Likewise, P values and sample sizes are reported in the figure and table legends or in dedicated tables.

## Declaration of generative AI and AI-assisted technologies in the manuscript preparation process

During the preparation of this work, the authors used Claude 4.6 to edit the manuscript for spelling, grammar and clarity. The authors reviewed and edited the output as needed and take full responsibility for the content of the manuscript.

## Acknowledgements

We thank the laboratories of C. Norden and M. Rocha-Martins for project discussions; João Coelho for experimental support; the Microscopy and Fish Facilities of the Gulbenkian Institute for Molecular Medicine (GIMM) and MPI for Molecular Biomedicine for technical support. C. Figueiredo is a member of the Integrative Biology and Biomedicine PhD programme and is supported by the Fundação para a Ciência e Tecnologia PhD fellowship (UI/BD/152255/2021). C. Norden was supported by the Fundação Calouste Gulbenkian-IGC, by the European Research Council (ERC) under the European Union’s Horizon 2020 research and innovation program (grant agreement no. 819046), and by the Fundação para a Ciência e a Tecnologia CEEC (2023.07063.CEECIND/CP2854/CT0002). G. Tellkamp is a member of the CiM-IMPRS Graduate School and is supported by the Max Planck Society. M. Rocha-Martins is supported by the MPI of Molecular Biomedicine and the Max Planck Society.

## Author contributions

C. Figueiredo: conceptualization, data curation, formal analysis, investigation, methodology, validation, visualization, writing (original draft). G. Tellkamp: investigation and methodology concerning p53 knockdown experiments. C. Norden: conceptualization, funding acquisition, project administration, resources, writing (original draft) and supervision. M. Rocha-Martins: conceptualization, funding acquisition, resources, investigation, methodology, writing (original draft), and supervision.

## Supplemental Material

**Figure S1.**
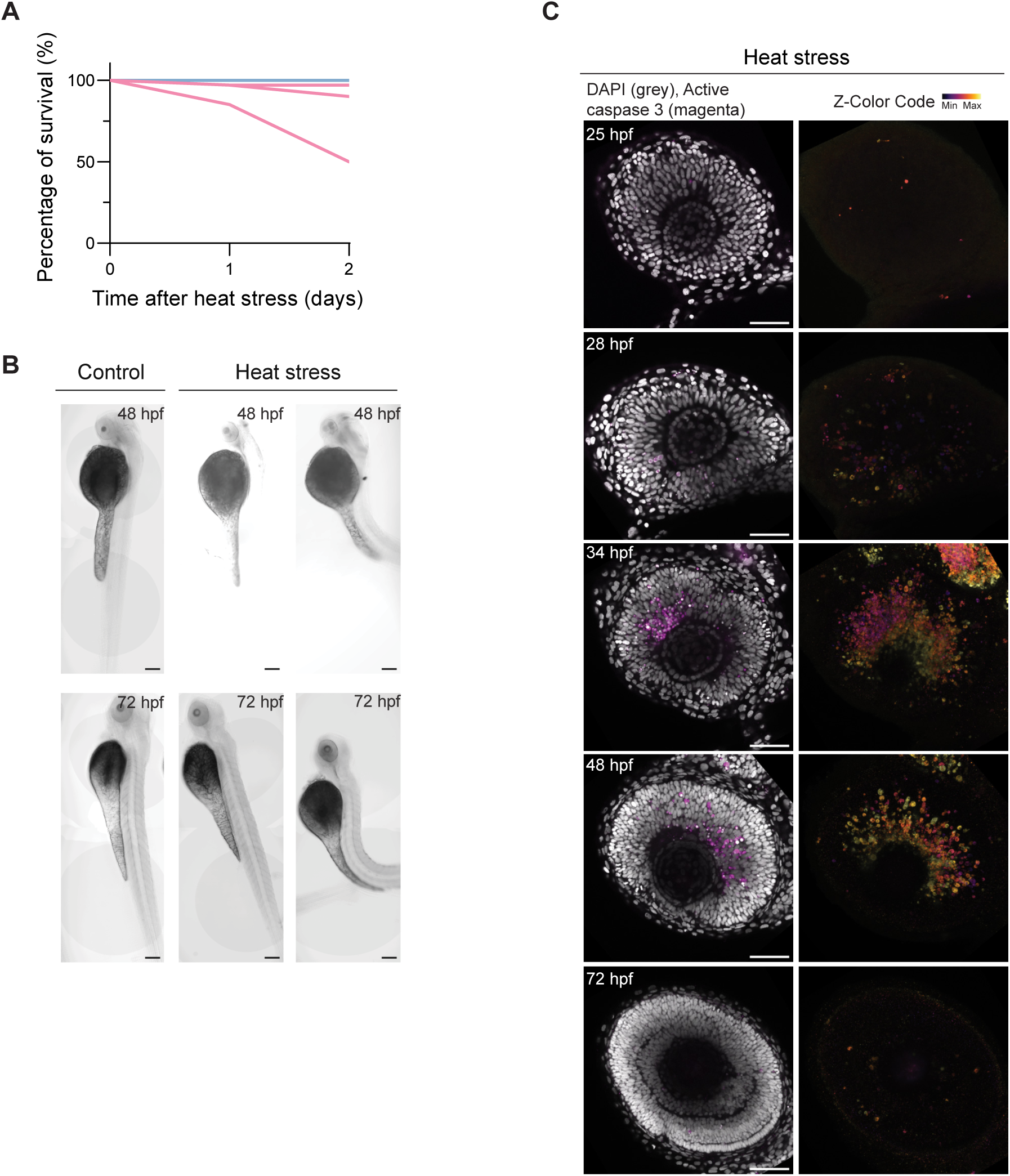
Heat stress induces transient apoptosis in the developing zebrafish retina. **(A)** Percentage of surviving embryos at 48 and 72 hpf in controls (blue; n=80) and after heat stress (pink; n=80) embryos (1 and 2 days after heat stress). Each line represents one replicate (3 replicates each for control and heat stress). **(B)** Brightfield images of control and heat-stressed embryos at 48 hpf and 72 hpf. **(C)** Spatial distribution of apoptotic cells in retinas of control (left) and heat-stressed (right) retinas at 0h (25 hpf), 4h (28 hpf), 10h (34 hpf), 24h (48 hpf) and 48h (72 hpf) after heat stress. DAPI (grey) labels nuclei. Active caspase 3 labels apoptotic cells; colour represents Z depth; lookup table shows minimum and maximum Z depths. Scale bars, 150 μm **(B)**, 50 μm **(C)**.

**Figure S2.**
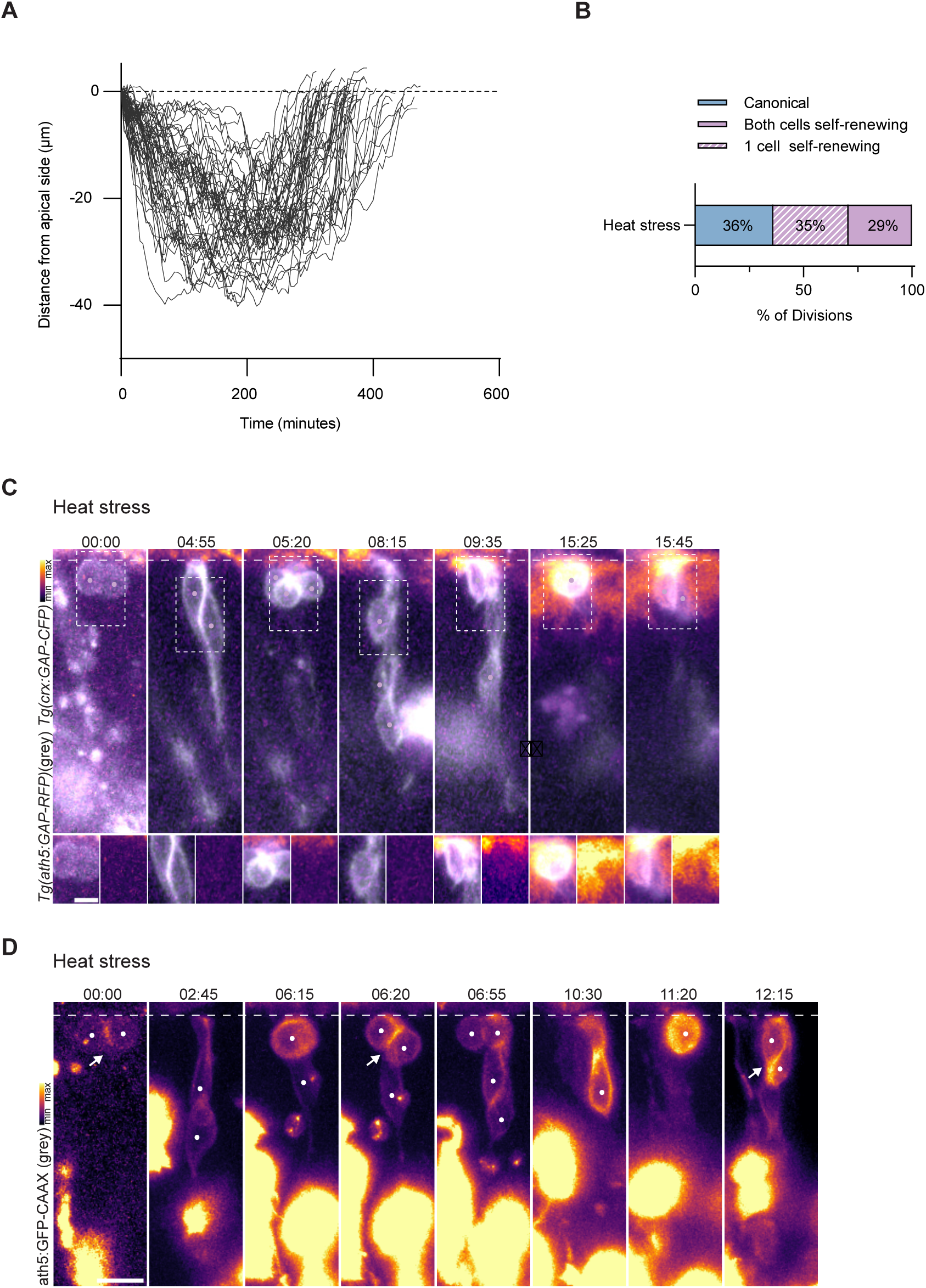
Ath5+ neurogenic progenitors undergo self-renewing divisions in heat stressed retinas. **(A)** Trajectories of self-renewing Ath5+ progenitors following heat stress (n=56 cells, N=14 embryos). Distance from the apical side is plotted over time. **(B)** Proportions of canonical and self-renewing divisions of one or both daughter cells of tracked Ath5+ progenitors in heat stressed retinas (n=72 cells, N=14 embryos). **(C)** Time series of a self-renewing Ath5+ neurogenic progenitor division in a heat-stressed embryo where progeny differentiates into PRpr. *Tg(ath5:GAP-RFP)* (grey) labels neurogenic progenitors and *Tg(crx:GAP-CFP)* labels PRprs. Dashed boxes indicate magnified regions shown below. In each timepoint, the left and right panels show merged RFP/CFP channels and the CFP channel, respectively. **(D)** Time series of a self-renewing Ath5+ neurogenic progenitor division in a heat-stressed embryo, in which the progeny undergoes a second self-renewing division. *Tg(ath5:GFP-CAAX)* labels Ath5+ neurogenic progenitors; colour represents fluorescence intensity. Lookup table shows minimum and maximum signal values. Arrow indicates cell divisions. **(C-D)** Dashed line indicates apical side. Time is displayed as hours:minutes. Scale bars, 10 μm **(C-D)** and 2 μm for magnified regions **(C)**.

**Figure S3.**
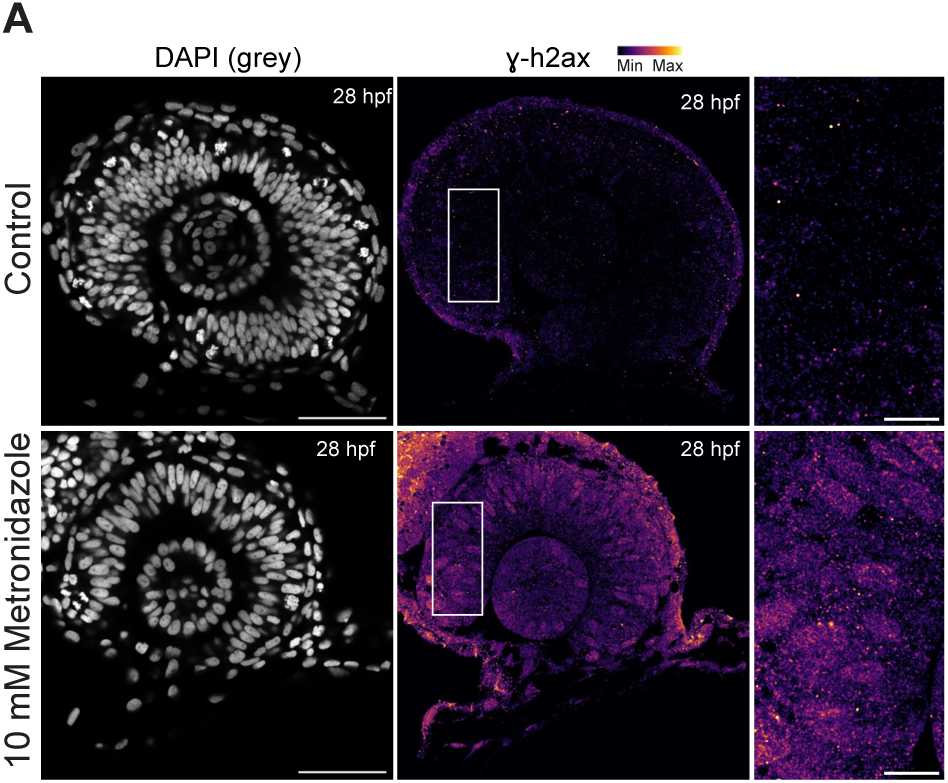
Chemogenetic ablation using the NTR/Mtz system induces DNA damage. (A) Retinas of control (top) and 10 mM Mtz-treated (bottom) CFP-NTR RNA-injected embryos. Mtz treatment occurred from 16-28 hpf. DAPI (grey) labels nuclei and γh2ax labels DNA double-strand breaks; colour represents fluorescence intensity. Lookup table shows minimum and maximum signal values. White boxes outline magnified regions (right). Scale bars, 50 μm, 10 μm for magnified regions.

**Figure S4.**
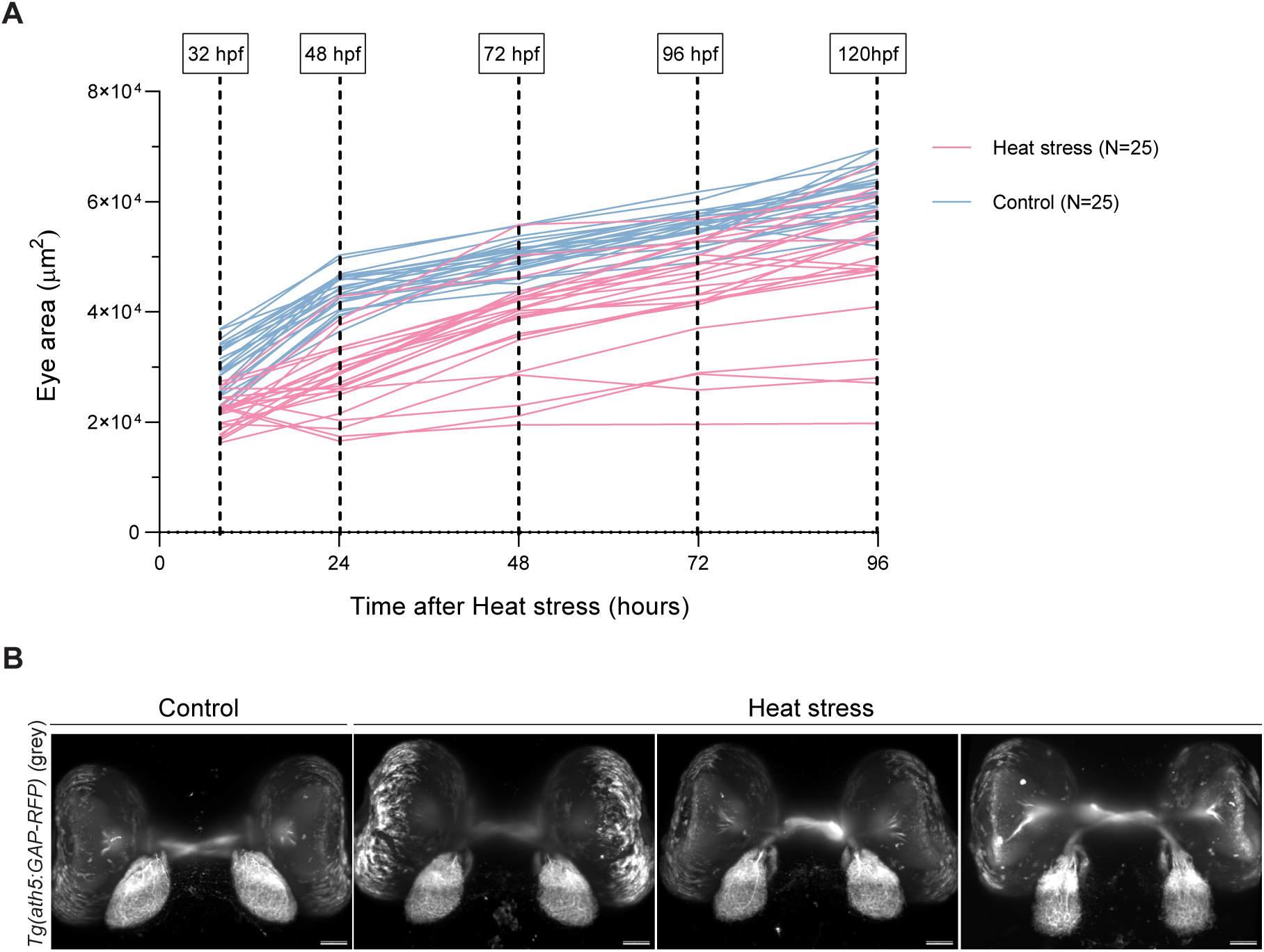
Heat-stressed retinas grow and form optic nerves that project to the brain. **(A)** Cohort of eye area measurements in control (blue, n=25) and heat-stressed (pink, n=25) embryos from 8 to 96 h after heat stress (32-120 hpf). Each line represents one embryo. Eye area is plotted over time. **(B)** Representative images of the optic nerves and neuronal projections to the tectum labelled by *Tg(ath5:GAP-RFP)* (grey) in control and heat-stressed embryos at 5 dpf. Scale bar, 50 μm.

## Video legends

**Video S1. Ath5+ neurogenic progenitors undergo self-renewing divisions in heat-stressed retinas, related to Figure 2**.

**Part 1.** Canonical division of Ath5+ progenitor in control retina, generating one basal neuron and one PRpr. ath5:GFP-CAAX (grey) labels early neurogenic progenitors and their progeny.

**Part 2-5** Effect of heat stress-induced apoptosis in Ath5+ progenitor divisions.

**Part 2.** Self-renewing division of Ath5+ progenitor. *Tg(ath5:GAP-RFP)* (grey) labels early neurogenic progenitors and their progeny.

**Part 3.** Canonical division of Ath5+ progenitor. *Tg(ath5:GAP-RFP)* (grey) labels early neurogenic progenitors and their progeny and *Tg(crx:GAP-CFP)* labels PRprs; colour represents fluorescence intensity. Lookup table shows min and max signal values.

**Part 4.** Self-renewing division of Ath5+ progenitor. One of the daughter cells of the self-renewing division undergoes a second, canonical, Ath5+ division giving rise to two CRX+ PRs and a basal neuron. ath5:GAP-RFP (grey) labels early neurogenic progenitors and their progeny and *Tg(crx:GAP-CFP)* labels PRprs; colour represents fluorescence intensity. Lookup table shows min and max signal values.

**Part 5.** Self-renewing division of Ath5+ progenitor in heat-stressed retina. Daughter cell of self-renewing division (red dot) undergoes a second self-renewing division. ath5:GFP-CAAX (grey) labels early neurogenic progenitors and their progeny.

Dots label tracked cells. Time is shown in hours:minutes. Scale bar is 10 μm.

**Video S2. Self-renewing divisions of Ath5+ neurogenic progenitors are a non-cell-autonomous response to apoptosis, related to Figure 3**.

**Part 1.** Self-renewing division of Ath5+ progenitor in CFP-NTR RNA-injected embryo. *Tg(ath5:GAP-RFP)* (grey) labels early neurogenic progenitors and their progeny.

**Part 2.** Self-renewing division of Ath5+ NTR-progenitor in rx2:CFP-NTR DNA-injected embryo. *Tg(ath5:GAP-RFP)* (grey) labels early neurogenic progenitors and their progeny.

**Part 3.** Self-renewing division of Ath5+ progenitor in control retina. Daughter cell of self-renewing division undergoes a second, canonical, Ath5+ division giving rise to two PRs and a basal neuron. ath5:GFP-CAAX (grey) and *Tg(ath5:GAP-RFP)* label early neurogenic progenitors and their progeny; colour represents fluorescence intensity. Lookup table shows min and max signal values.

Dots label tracked cells. Time is shown in hours:minutes. Scale bar is 10 μm.

## References

1. Hamdoun, A., and Epel, D. (2007). Embryo stability and vulnerability in an always changing world. Proc. Natl. Acad. Sci. U. S. A. 104, 1745–1750. 10.1073/pnas.0610108104.

2. Gilbert, S.F. (2001). Ecological Developmental Biology: Developmental Biology Meets the Real World. Dev. Biol. 233, 1–12. 10.1006/dbio.2001.0210.

3. Waddington, C.H. (1942). Canalization of Development and the Inheritance of Acquired Characters. Nature 150, 563–565. 10.1038/150563a0.

4. Wong, M., and Gilmour, D. (2020). Getting back on track: exploiting canalization to uncover the mechanisms of developmental robustness. Curr. Opin. Genet. Dev. 63, 53– 60. 10.1016/j.gde.2020.04.001.

5. González-Blanco, A., Acuña-Higaki, A., Boettger, D., Joy, J., and Milán, M. (2026). Proteostasis failure and mitochondrial dysfunction contribute to chromosomal instability-induced microcephaly. Nat. Commun. 10.1038/s41467-026-70521-0.

6. Marthiens, V., Rujano, M.A., Pennetier, C., Tessier, S., Paul-Gilloteaux, P., and Basto, R. (2013). Centrosome amplification causes microcephaly. Nat. Cell Biol. 15, 731–740. 10.1038/ncb2746.

7. Miao, H., Wu, H., Zhu, Y., Kong, L., Yu, X., Zeng, Q., Chen, Y., Zhang, Q., Guo, P., and Wang, D. (2021). Congenital anomalies associated with ambient temperature variability during fetal organogenesis period of pregnancy: Evidence from 4.78 million births. Sci. Total Environ. 798, 149305. 10.1016/j.scitotenv.2021.149305.

8. Dorrity, M.W., Saunders, L.M., Duran, M., Srivatsan, S.R., Barkan, E., Jackson, D.L., Sattler, S.M., Ewing, B., Queitsch, C., Shendure, J., et al. (2023). Proteostasis governs differential temperature sensitivity across embryonic cell types. Cell 186, 5015–5027.e12. 10.1016/j.cell.2023.10.013.

9. Menon, T., and Nair, S. (2018). Transient window of resilience during early development minimizes teratogenic effects of heat in zebrafish embryos. Dev. Dyn. 247, 992–1004. 10.1002/dvdy.24640.

10. Wang, R., Zhang, H., Du, J., and Xu, J. (2019). Heat resilience in embryonic zebrafish revealed using an in vivo stress granule reporter. J. Cell Sci. 132, jcs234807. 10.1242/jcs.234807.

11. Hughes, S., Vrinds, I., de Roo, J., Francke, C., Shimeld, S.M., Woollard, A., and Sato, A. (2019). DnaJ chaperones contribute to canalization. J. Exp. Zool. Part Ecol. Integr. Physiol. 331, 201–212. 10.1002/jez.2254.

12. McClellan, A.J., Tam, S., Kaganovich, D., and Frydman, J. (2005). Protein quality control: chaperones culling corrupt conformations. Nat. Cell Biol. 7, 736–741. 10.1038/ncb0805-736.

13. Saibil, H. (2013). Chaperone machines for protein folding, unfolding and disaggregation. Nat. Rev. Mol. Cell Biol. 14, 630–642. 10.1038/nrm3658.

14. Cannavò, E., Khoueiry, P., Garfield, D.A., Geeleher, P., Zichner, T., Gustafson, E.H., Ciglar, L., Korbel, J.O., and Furlong, E.E.M. (2016). Shadow Enhancers Are Pervasive Features of Developmental Regulatory Networks. Curr. Biol. 26, 38–51. 10.1016/j.cub.2015.11.034.

15. Sokolowski, T.R., Erdmann, T., and ten Wolde, P.R. (2012). Mutual Repression Enhances the Steepness and Precision of Gene Expression Boundaries. PLoS Comput. Biol. 8, e1002654. 10.1371/journal.pcbi.1002654.

16. Akieda, Y., Ogamino, S., Furuie, H., Ishitani, S., Akiyoshi, R., Nogami, J., Masuda, T., Shimizu, N., Ohkawa, Y., and Ishitani, T. (2019). Cell competition corrects noisy Wnt morphogen gradients to achieve robust patterning in the zebrafish embryo. Nat. Commun. 10, 4710. 10.1038/s41467-019-12609-4.

17. Jelier, R., Kruger, A., Swoger, J., Zimmermann, T., and Lehner, B. (2016). Compensatory Cell Movements Confer Robustness to Mechanical Deformation during Embryonic Development. Cell Syst. 3, 160–171. 10.1016/j.cels.2016.07.005.

18. Pinet, K., Deolankar, M., Leung, B., and McLaughlin, K.A. (2019). Adaptive correction of craniofacial defects in pre-metamorphic Xenopus laevis tadpoles involves thyroid hormone-independent tissue remodeling. Development 146, dev175893. 10.1242/dev.175893.

19. Huh, J.R., Guo, M., and Hay, B.A. (2004). Compensatory Proliferation Induced by Cell Death in the Drosophila Wing Disc Requires Activity of the Apical Cell Death Caspase Dronc in a Nonapoptotic Role. Curr. Biol. 14, 1262–1266. 10.1016/j.cub.2004.06.015.

20. Kondo, S., Senoo-Matsuda, N., Hiromi, Y., and Miura, M. (2006). DRONC Coordinates Cell Death and Compensatory Proliferation. Mol. Cell. Biol. 26, 7258–7268. 10.1128/MCB.00183-06.

21. Milán, M., Campuzano, S., and García-Bellido, A. (1997). Developmental parameters of cell death in the wing disc of *Drosophila*. Proc. Natl. Acad. Sci. 94, 5691–5696. 10.1073/pnas.94.11.5691.

22. Freret-Hodara, B., Cui, Y., Griveau, A., Vigier, L., Arai, Y., Touboul, J., and Pierani, A. (2017). Enhanced Abventricular Proliferation Compensates Cell Death in the Embryonic Cerebral Cortex. Cereb. Cortex N. Y. N 1991 *27*, 4701–4718. 10.1093/cercor/bhw264.

23. Fan, Y., and Bergmann, A. (2008). Distinct Mechanisms of Apoptosis-Induced Compensatory Proliferation in Proliferating and Differentiating Tissues in the Drosophila Eye. Dev. Cell 14, 399–410. 10.1016/j.devcel.2008.01.003.

24. Ryoo, H.D., Gorenc, T., and Steller, H. (2004). Apoptotic Cells Can Induce Compensatory Cell Proliferation through the JNK and the Wingless Signaling Pathways. Dev. Cell 7, 491– 501. 10.1016/j.devcel.2004.08.019.

25. Buss, R.R., Sun, W., and Oppenheim, R.W. (2006). ADAPTIVE ROLES OF PROGRAMMED CELL DEATH DURING NERVOUS SYSTEM DEVELOPMENT. Annu. Rev. Neurosci. 29, 1–35. 10.1146/annurev.neuro.29.051605.112800.

26. Homem, C.C.F., Repic, M., and Knoblich, J.A. (2015). Proliferation control in neural stem and progenitor cells. Nat. Rev. Neurosci. 16, 647–659. 10.1038/nrn4021.

27. Biehlmaier, O., Neuhauss, S.C., and Kohler, K. (2001). Onset and time course of apoptosis in the developing zebrafish retina. Cell Tissue Res. 306, 199–207. 10.1007/s004410100447.

28. Vecino, E., Hernandez, M., and Garcia, M. (2004). Cell death in the developing vertebrate retina. Int. J. Dev. Biol. 48, 965–974. 10.1387/ijdb.041891ev.

29. Yoshida, H., Kong, Y.-Y., Yoshida, R., Elia, A.J., Hakem, A., Hakem, R., Penninger, J.M., and Mak, T.W. (1998). Apaf1 Is Required for Mitochondrial Pathways of Apoptosis and Brain Development. Cell 94, 739–750. 10.1016/S0092-8674(00)81733-X.

30. Garcez, P.P., Loiola, E.C., Madeiro da Costa, R., Higa, L.M., Trindade, P., Delvecchio, R., Nascimento, J.M., Brindeiro, R., Tanuri, A., and Rehen, S.K. (2016). Zika virus impairs growth in human neurospheres and brain organoids. Science 352, 816–818. 10.1126/science.aaf6116.

31. Matos-Rodrigues, G.E., Tan, P.B., Rocha-Martins, M., Charlier, C.F., Gomes, A.L., Cabral-Miranda, F., Grigaravicius, P., Hofmann, T.G., Frappart, P.-O., and Martins, R.A.P. (2020). Progenitor death drives retinal dysplasia and neuronal degeneration in a mouse model of ATRIP-Seckel syndrome. Dis. Model. Mech. 13, dmm045807. 10.1242/dmm.045807.

32. European Platform on Rare Disease Registration https://eu-rd-platform.jrc.ec.europa.eu.

33. Lim, N.R., Shohayeb, B., Zaytseva, O., Mitchell, N., Millard, S.S., Ng, D.C.H., and Quinn, L.M. (2017). Glial-Specific Functions of Microcephaly Protein WDR62 and Interaction with the Mitotic Kinase AURKA Are Essential for Drosophila Brain Growth. Stem Cell Rep. 9, 32–41. 10.1016/j.stemcr.2017.05.015.

34. van Rahden, V.A., Fernandez-Vizarra, E., Alawi, M., Brand, K., Fellmann, F., Horn, D., Zeviani, M., and Kutsche, K. (2015). Mutations in NDUFB11, Encoding a Complex I Component of the Mitochondrial Respiratory Chain, Cause Microphthalmia with Linear Skin Defects Syndrome. Am. J. Hum. Genet. 96, 640–650. 10.1016/j.ajhg.2015.02.002.

35. Diaz, C., and Glover, J.C. (1996). Appropriate pattern formation following regulative regeneration in the hindbrain neural tube. Development 122, 3095–3105. 10.1242/dev.122.10.3095.

36. Halasi, G., Søviknes, A.M., Sigurjonsson, O., and Glover, J.C. (2012). Proliferation and recapitulation of developmental patterning associated with regulative regeneration of the spinal cord neural tube. Dev. Biol. 365, 118–132. 10.1016/j.ydbio.2012.02.012.

37. Lindenhofer, D., Haendeler, S., Esk, C., Littleboy, J.B., Brunet Avalos, C., Naas, J., Pflug, F.G., van de Ven, E.G.P., Reumann, D., Baffet, A.D., et al. (2024). Cerebral organoids display dynamic clonal growth and tunable tissue replenishment. Nat. Cell Biol. 26, 710–718. 10.1038/s41556-024-01412-z.

38. Dzafic, E., Strzyz, P.J., Wilsch-Bräuninger, M., and Norden, C. (2015). Centriole Amplification in Zebrafish Affects Proliferation and Survival but Not Differentiation of Neural Progenitor Cells. Cell Rep. 13, 168–182. 10.1016/j.celrep.2015.08.062.

39. Boije, H., Rulands, S., Dudczig, S., Simons, B.D., and Harris, W.A. (2015). The Independent Probabilistic Firing of Transcription Factors: A Paradigm for Clonal Variability in the Zebrafish Retina. Dev. Cell 34, 532–543. 10.1016/j.devcel.2015.08.011.

40. He, J., Zhang, G., Almeida, A.D., Cayouette, M., Simons, B.D., and Harris, W.A. (2012). How Variable Clones Build an Invariant Retina. Neuron 75, 786–798. 10.1016/j.neuron.2012.06.033.

41. Llorca, A., Ciceri, G., Beattie, R., Wong, F.K., Diana, G., Serafeimidou-Pouliou, E., Fernández-Otero, M., Streicher, C., Arnold, S.J., Meyer, M., et al. (2019). A stochastic framework of neurogenesis underlies the assembly of neocortical cytoarchitecture. eLife 8, e51381. 10.7554/eLife.51381.

42. Matejčić, M., Salbreux, G., and Norden, C. (2018). A non-cell-autonomous actin redistribution enables isotropic retinal growth. PLOS Biol. 16, e2006018. 10.1371/journal.pbio.2006018.

43. Nerli, E., Rocha-Martins, M., and Norden, C. (2020). Asymmetric neurogenic commitment of retinal progenitors involves Notch through the endocytic pathway. eLife 9, e60462. 10.7554/eLife.60462.

44. Nerli, E., Kretzschmar, J., Bianucci, T., Rocha-Martins, M., Zechner, C., and Norden, C. (2023). Deterministic and probabilistic fate decisions co-exist in a single retinal lineage. EMBO J. n/a, e112657. 10.15252/embj.2022112657.

45. Weber, I.P., Ramos, A.P., Strzyz, P.J., Leung, L.C., Young, S., and Norden, C. (2014). Mitotic Position and Morphology of Committed Precursor Cells in the Zebrafish Retina Adapt to Architectural Changes upon Tissue Maturation. Cell Rep. 7, 386–397. 10.1016/j.celrep.2014.03.014.

46. Edwards, M.J. (1998). Apoptosis, the heat shock response, hyperthermia, birth defects, disease and cancer. Where are the common links? Cell Stress Chaperones 3, 213–220. 10.1379/1466-1268(1998)003%3C0213:athsrh%3E2.3.co;2.

47. Khan, V.R., and Brown, I.R. (2002). The effect of hyperthermia on the induction of cell death in brain, testis, and thymus of the adult and developing rat. Cell Stress Chaperones 7, 73–90. 10.1379/1466-1268(2002)007%3C0073:teohot%3E2.0.co;2.

48. Sanger, T.J., Harding, L., Kyrkos, J., Turnquist, A.J., Epperlein, L., Nunez, S.A., Lachance, D., Dhindsa, S., Stroud, J.T., Diaz, R.E., Jr, et al. (2021). Environmental Thermal Stress Induces Neuronal Cell Death and Developmental Malformations in Reptiles. Integr. Org. Biol. 3, obab033. 10.1093/iob/obab033.

49. Yabu, T., Todoriki, S., and Yamashita, M. (2001). Stress-induced apoptosis by heat shock, UV and γ-ray irradiation in zebrafish embryos detected by increased caspase activity and whole-mount TUNEL staining. Fish. Sci. 67, 333–340. 10.1046/j.1444-2906.2001.00233.x.

50. Bellmann, K., Charette, S.J., Nadeau, P.J., Poirier, D.J., Loranger, A., and Landry, J. (2010). The mechanism whereby heat shock induces apoptosis depends on the innate sensitivity of cells to stress. Cell Stress Chaperones 15, 101–113. 10.1007/s12192-009-0126-9.

51. Takahashi, A., Matsumoto, H., Nagayama, K., Kitano, M., Hirose, S., Tanaka, H., Mori, E., Yamakawa, N., Yasumoto, J., Yuki, K., et al. (2004). Evidence for the Involvement of Double-Strand Breaks in Heat-Induced Cell Killing. Cancer Res. 64, 8839–8845. 10.1158/0008-5472.CAN-04-1876.

52. Warters, R.L., and Brizgys, L.M. (1987). Apurinic site induction in the DNA of cells heated at hyperthermic temperatures. J. Cell. Physiol. 133, 144–150. 10.1002/jcp.1041330118.

53. Robu, M.E., Larson, J.D., Nasevicius, A., Beiraghi, S., Brenner, C., Farber, S.A., and Ekker, S.C. (2007). p53 Activation by Knockdown Technologies. PLOS Genet. 3, e78. 10.1371/journal.pgen.0030078.

54. Ho H’ng, C., Amarasinghe, S.L., Zhang, B., Chang, H., Qu, X., Powell, D.R., and Rosello-Diez, A. (2024). Compensatory growth and recovery of cartilage cytoarchitecture after transient cell death in fetal mouse limbs. Nat. Commun. 15, 2940. 10.1038/s41467-024-47311-7.

55. Young, R.M., Hawkins, T.A., Cavodeassi, F., Stickney, H.L., Schwarz, Q., Lawrence, L.M., Wierzbicki, C., Cheng, B.Y., Luo, J., Ambrosio, E.M., et al. (2019). Compensatory growth renders Tcf7l1a dispensable for eye formation despite its requirement in eye field specification. eLife 8, e40093. 10.7554/eLife.40093.

56. Brown, N.L., Kanekar, S., Vetter, M.L., Tucker, P.K., Gemza, D.L., and Glaser, T. (1998). Math5 encodes a murine basic helix-loop-helix transcription factor expressed during early stages of retinal neurogenesis. Development 125, 4821–4833. 10.1242/dev.125.23.4821.

57. Poggi, L., Vitorino, M., Masai, I., and Harris, W.A. (2005). Influences on neural lineage and mode of division in the zebrafish retina in vivo. J. Cell Biol. 171, 991–999. 10.1083/jcb.200509098.

58. Zolessi, F.R., Poggi, L., Wilkinson, C.J., Chien, C.-B., and Harris, W.A. (2006). Polarization and orientation of retinal ganglion cells in vivo. Neural Develop. 1, 2. 10.1186/1749-8104-1-2.

59. Icha, J., Weber, M., Waters, J.C., and Norden, C. (2017). Phototoxicity in live fluorescence microscopy, and how to avoid it. BioEssays 39, 1700003. 10.1002/bies.201700003.

60. Rocha-Martins, M., Nerli, E., Kretzschmar, J., Weigert, M., Icha, J., Myers, E.W., and Norden, C. (2023). Neuronal migration prevents spatial competition in retinal morphogenesis. Nature 620, 615–624. 10.1038/s41586-023-06392-y.

61. Nerli, E. (2023). Investigating progenitor cell lineages and their regulation during zebrafish retinal neurogenesis.

62. Vitorino, M., Jusuf, P.R., Maurus, D., Kimura, Y., Higashijima, S., and Harris, W.A. (2009). Vsx2 in the zebrafish retina: restricted lineages through derepression. Neural Develop. 4, 14. 10.1186/1749-8104-4-14.

63. Clark, A.J., Iwobi, M., Cui, W., Crompton, M., Harold, G., Hobbs, S., Kamalati, T., Knox, R., Neil, C., Yull, F., et al. (1997). Selective cell ablation in transgenic mice expressing E. coli nitroreductase. Gene Ther. 4, 101–110. 10.1038/sj.gt.3300367.

64. Curado, S., Stainier, D.Y.R., and Anderson, R.M. (2008). Nitroreductase-mediated cell/tissue ablation in zebrafish: a spatially and temporally controlled ablation method with applications in developmental and regeneration studies. Nat. Protoc. 3, 948–954. 10.1038/nprot.2008.58.

65. Almeida, A.D., Boije, H., Chow, R.W., He, J., Tham, J., Suzuki, S.C., and Harris, W.A. (2014). Spectrum of Fates: a new approach to the study of the developing zebrafish retina. Development 141, 1971–1980. 10.1242/dev.104760.

66. Prakash, B.A., and Toro, C.P. (2019). Modulating the Zebrafish Camouflage Response Pathway to Illustrate Neuroendocrine Control Over a Robust and Quantifiable Behavior. J. Undergrad. Neurosci. Educ. 18, A57–A64.

67. Slanchev, K., Laurell, E., Arnold-Ammer, I., Kuehn, E., Pandiarajan, U., and Baier, H. (2025). Molecular delineation of a retina-dependent photoneuroendocrine pathway in zebrafish. Curr. Biol. 10.1016/j.cub.2025.10.025.

68. Gomes, F.L.A.F., Zhang, G., Carbonell, F., Correa, J.A., Harris, W.A., Simons, B.D., and Cayouette, M. (2011). Reconstruction of rat retinal progenitor cell lineages in vitro reveals a surprising degree of stochasticity in cell fate decisions. Development 138, 227–235. 10.1242/dev.059683.

69. Iribarne, M., Hyde, D.R., and Masai, I. (2019). TNFα Induces Müller Glia to Transition From Non-proliferative Gliosis to a Regenerative Response in Mutant Zebrafish Presenting Chronic Photoreceptor Degeneration. Front. Cell Dev. Biol. 7. 10.3389/fcell.2019.00296.

70. Nelson, C.M., Ackerman, K.M., O’Hayer, P., Bailey, T.J., Gorsuch, R.A., and Hyde, D.R. (2013). Tumor Necrosis Factor-Alpha Is Produced by Dying Retinal Neurons and Is Required for Müller Glia Proliferation during Zebrafish Retinal Regeneration. J. Neurosci. 33, 6524–6539. 10.1523/JNEUROSCI.3838-12.2013.

71. Kawaue, T., Yow, I., Pan, Y., Le, A.P., Lou, Y., Loberas, M., Shagirov, M., Teng, X., Prost, J., Hiraiwa, T., et al. (2023). Inhomogeneous mechanotransduction defines the spatial pattern of apoptosis-induced compensatory proliferation. Dev. Cell 58, 267–277.e5. 10.1016/j.devcel.2023.01.005.

72. Cavone, L., McCann, T., Drake, L.K., Aguzzi, E.A., Oprişoreanu, A.-M., Pedersen, E., Sandi, S., Selvarajah, J., Tsarouchas, T.M., Wehner, D., et al. (2021). A unique macrophage subpopulation signals directly to progenitor cells to promote regenerative neurogenesis in the zebrafish spinal cord. Dev. Cell 56, 1617–1630.e6. 10.1016/j.devcel.2021.04.031.

73. Fogarty, C.E., Diwanji, N., Lindblad, J.L., Tare, M., Amcheslavsky, A., Makhijani, K., Brückner, K., Fan, Y., and Bergmann, A. (2016). Extracellular Reactive Oxygen Species Drive Apoptosis-Induced Proliferation via Drosophila Macrophages. Curr. Biol. 26, 575– 584. 10.1016/j.cub.2015.12.064.

74. Mellén, M.A., de la Rosa, E.J., and Boya, P. (2008). The autophagic machinery is necessary for removal of cell corpses from the developing retinal neuroepithelium. Cell Death Differ. 15, 1279–1290. 10.1038/cdd.2008.40.

75. Wang, M., Du, L., Lee, A.C., Li, Y., Qin, H., and He, J. (2020). Different lineage contexts direct common pro-neural factors to specify distinct retinal cell subtypes. J. Cell Biol. 219, e202003026. 10.1083/jcb.202003026.

76. Gibson, G., and Dworkin, I. (2004). Uncovering cryptic genetic variation. Nat. Rev. Genet. 5, 681–690. 10.1038/nrg1426.

77. Hayden, E.J., Ferrada, E., and Wagner, A. (2011). Cryptic genetic variation promotes rapid evolutionary adaptation in an RNA enzyme. Nature 474, 92–95. 10.1038/nature10083.

78. Zheng, J., Payne, J.L., and Wagner, A. (2019). Cryptic genetic variation accelerates evolution by opening access to diverse adaptive peaks. Science 365, 347–353. 10.1126/science.aax1837.

79. Icha, J., Kunath, C., Rocha-Martins, M., and Norden, C. (2016). Independent modes of ganglion cell translocation ensure correct lamination of the zebrafish retina. J. Cell Biol. 215, 259–275. 10.1083/jcb.201604095.

80. Kay, J.N., Link, B.A., and Baier, H. (2005). Staggered cell-intrinsic timing of *ath5* expression underlies the wave of ganglion cell neurogenesis in the zebrafish retina. Development 132, 2573–2585. 10.1242/dev.01831.

81. Kimmel, C.B., Ballard, W.W., Kimmel, S.R., Ullmann, B., and Schilling, T.F. (1995). Stages of embryonic development of the zebrafish. Dev. Dyn. 203, 253–310. 10.1002/aja.1002030302.

82. Kwan, K.M., Fujimoto, E., Grabher, C., Mangum, B.D., Hardy, M.E., Campbell, D.S., Parant, J.M., Yost, H.J., Kanki, J.P., and Chien, C.-B. (2007). The Tol2kit: A multisite gateway-based construction kit for Tol2 transposon transgenesis constructs. Dev. Dyn. 236, 3088–3099. 10.1002/dvdy.21343.

83. Martinez-Morales, J.R., Rembold, M., Greger, K., Simpson, J.C., Brown, K.E., Quiring, R., Pepperkok, R., Martin-Bermudo, M.D., Himmelbauer, H., and Wittbrodt, J. (2009). ojoplano-mediated basal constriction is essential for optic cup morphogenesis. Development 136, 2165–2175. 10.1242/dev.033563.

84. Villefranc, J.A., Amigo, J., and Lawson, N.D. (2007). Gateway compatible vectors for analysis of gene function in the zebrafish. Dev. Dyn. 236, 3077–3087. 10.1002/dvdy.21354.

85. Icha, J., Schmied, C., Sidhaye, J., Tomancak, P., Preibisch, S., and Norden, C. (2016). Using Light Sheet Fluorescence Microscopy to Image Zebrafish Eye Development. J. Vis. Exp. JoVE, 53966. 10.3791/53966.

86. Soans, K.G., Ramos, A.P., Sidhaye, J., Krishna, A., Solomatina, A., Hoffmann, K.B., Schlüßler, R., Guck, J., Sbalzarini, I.F., Modes, C.D., et al. (2022). Collective cell migration during optic cup formation features changing cell-matrix interactions linked to matrix topology. Curr. Biol. 32, 4817–4831.e9. 10.1016/j.cub.2022.09.034.

87. Muto, A., Orger, M.B., Wehman, A.M., Smear, M.C., Kay, J.N., Page-McCaw, P.S., Gahtan, E., Xiao, T., Nevin, L.M., Gosse, N.J., et al. (2005). Forward Genetic Analysis of Visual Behavior in Zebrafish. PLOS Genet. 1, e66. 10.1371/journal.pgen.0010066.

88. Gorelik, R., and Gautreau, A. (2014). Quantitative and unbiased analysis of directional persistence in cell migration. Nat. Protoc. 9, 1931–1943. 10.1038/nprot.2014.131.

